# The dual coding of a single sex pheromone receptor in regulating the mating behavior of Asian Honeybee *Apis cerana*

**DOI:** 10.1101/2023.12.03.569826

**Authors:** Haoqin Ke, Jonathan D. Bohbot, Yongjuan Chi, Shiwen Duan, Xiaomei Ma, Bingzhong Ren, Yinliang Wang

## Abstract

In honeybee society, a virgin queen usually mates only once with several drones before founding a colony. For the rest of her prolific life, she will not engage in subsequent mating events. In the Asian honeybee *Apis cerana*, the mechanisms controlling this reproductive strategy involves the queen-released mandibular pheromone (QMP). This pheromone blend regulates the physiology and reproductive behavior of workers and drones. Its main component, 9-oxo-(*E*)-2-decenoic acid (9-ODA), acts as a sex pheromone and attracts drones. However, how the QMP prevents additional mating later in the queen’s life remains elusive. Using behavioral and chemical analysis, we show that the QMP component methyl *p*-hydroxybenzoate (HOB) released by mated queens inhibits male attraction to 9-ODA. Furthermore, *in vivo* electroantennogram and single sensillum recording data indicated that HOB alone significantly reduces the spontaneous spike activity of 9-ODA-sensitive olfactory receptor neurons (ORNs). To explore the molecular mechanism underlying this inverse effect, we conducted qPCR and in situ hybridization assays. The results indicated that *AcerOr11* is specifically expressed in sensilla placodea from the drone’s antennae, which are the sensilla that narrowly respond to both 9-ODA and HOB. We then cloned and expressed *AcerOr11* in a Xenopus oocyte expression system, where *AcerOr11* induced robust inward (regular) currents in response to 9-ODA. Intriguingly, HOB induced inverse currents in a dose-dependent manner. This suggests that HOB may act as an inverse agonist against *AcerOr11*, providing additional odor-coding information. Based on these findings, we propose a model in which *AcerOr11* function as a dual modulators in regulating the mating behavior of *A. cerana*. The inverse agonist, HOB, can help manage the population dynamics of honeybees in apiculture.

## Introduction

Population dynamics in insects are influenced by various environmental factors, including temperature, humidity, light, vegetative biodiversity on diets, etc. ^1^. In eusocial insects, such as honeybees, population dynamics become more complex due to the interdependence of individuals within the colony, and their social behavior is tightly regulated ^2^. Besides, environmental factors and intraspecific chemical signals play a crucial role in the growth and stability of the honeybee colony by regulating reproduction. For example, fertile *Apis mellifera* workers are identified by the esters content in Dufour’s gland secretions, while the worker-laid eggs are attacked by other workers ^3^. Moreover, brood signals such as aliphatic esters and E-*β*-ocimene, produced by *A. mellifera* larvae, not only elicit brood care behaviors but also suppress worker reproduction ^4^.

Among these chemical signals, the queen mandibular pheromone (QMP) plays a key role in regulating colony reproduction. QMP directly triggers the mating behavior of drones and maintains the reproductive monopoly of the queen, thereby moderating population dynamics in honeybees. QMP provides information about the mating status of females and attracts drones, while also inhibiting the development of worker ovaries by changing relevant gene expression ^5^. Several studies have focused on *A. mellifera* QMP, which consists of four main components: 9-oxo-(*E*)-2-decenoic acid (9-ODA), (*R*, *S*)-9-hydroxy-(*E*)-2-decenoic acid (9-HDA), methyl *p*-hydroxybenzoate (HOB), and 4-hydroxy-3-methoxyphenyl-ethanol (HVA) ^6^. To date, only 9-ODA has been shown to have an attractive effect on drones ^5^. In *A. cerana*, 9-ODA, 9-HDA and HOB are also the dominant component of QMP, but HVA is absent. Behavioral evidence suggests that the QMP mixture lacking HVA in *A. cerana* is still sufficient to elicit retinue behavior in workers ^6^, indicating a similar function of QMP in *A. cerana* to that in *A. mellifera*. However, the role of HOB remains poorly understood, as single secondary QMP components are not attractive. Until this study, the mechanism by which HOB alone regulates mating behavior in *A. cerana* has remained unknown.

Olfactory sensing of QMP has been extensively studied in *A. mellifera*. At the peripheral olfactory system level, the primary QMP component, 9-ODA, is detected by the placoid sensilla located on the drone antenna, which expresses the *Apis mellifera* odorant receptor 11 gene (*AmelOr11*). When expressed in the *Xenopus laevis* system, *AmelOr11* exhibits a narrow response to 9-ODA and does not respond to other QMP components ^7^. This suggests the involvement of other olfactory proteins in detecting QMP secondary components. The predominant role of 9-ODA in mating is further supported by calcium imaging experiments at the antennal lobe level, where the *A. mellifera* macroglomerulus MG2 in drones is activated by 9-ODA ^8^. Although *A. mellifera* and *A. cerana* share overall similar morphology and social behavior, they differ in certain aspects of their olfactory systems, such as QMP composition, the number of odorant receptors (*Ors*), and antennal lobe topology ^9^. The QMP olfactory sensing in *A. cerana* has received less attention compared to *A. mellifera*. In *A. cerana*, odorant binding protein 11 (AcerOBP11) demonstrates strong binding affinities for both 9-ODA and HOB ^10^. However, the specific odorant receptors (ORs) responsible for detecting 9-ODA and HOB in *A. cerana* remain unclear.

In this study, we found that HOB, released only by mated queens, significantly reduce the attraction of drones to 9-ODA. This inverse effect of HOB was further validated by *in vivo* electrophysiological assays, electroantennogram (EAG), and single sensillum recording (SSR). Lastly, we uncovered that, similar to the *A. mellifera* orthologs, *AcerOr11* is robustly activated by 9-ODA, moreover, the secondary component HOB elicited reverse current fluxes, implying the existence of a dual coding mechanism at the QMP receptor, *AcerOr11*. This study aimed to explore how QMP regulates reproduction at the olfactory sensing level and modulates the population dynamics of *A. cerana*.

## Materials and methods

### Honeybees

Honeybees (*A. cerana*) used in this study were provided by the Jilin Provincial Institute of Apicultural Sciences (JLAS), China. The bee colony was originally collected from Dunhua, China (43° 51′ 46′′ N, 128° 20′ 30′′ E) and has been reared since 2020 in the conservation area of JLAS in a natural environment. Before beginning the experiments, the honeybees were cultured in an artificial incubator (Boxun, China) at 30 ℃, 70% humidity, and in a 16-h photoperiod. Adult bees were fed with a 10% sucrose solution.

### Gas chromatographic-mass spectrometry (GC-MS) analysis

Heads from the virgin queens, mated queens (mated on day 6 or 7), drones, and workers were collected from 12- to 15-day-old honeybees. The compounds of the mandibular gland were extracted by placing the heads in 200 μL dichloromethane for at least 24 h. The extracts then dried under a stream of nitrogen, and the remainders were dissolved in 20 μL internal standard solution (octanoic acid and tetradecane in dichloromethane) and 20 μL N, O-Bis(trimethylsilyl) trifluoroacetamide (BSTFA). The mandibular gland pheromone mixes were separated by a GC-MS system (Agilent 6890N/5973I, Thermo Fisher Scientific, USA), which was equipped with an HP-INNOWax capillary column, in the split-less mode on a methyl silicone-coated fused silica column (HP - 1MS, 25 m × 0.20 mm × 0.33 µm). Helium gas was used as a carrier gas at a constant flow rate of 1 mL/min. The oven temperature was set at 100 °C for 2 min and then increased to 250 °C at a rate of 10 °C per minute. The final temperature was maintained for 10 min. The compounds were identified by comparing their retention times with the known reference compounds. 9-ODA and HOB were further quantified by injecting corresponding standard compounds.

### Behavioral assay

To explore the biological effect of 9-ODA and HOB on drones, we designed a binary-choice Y-tube olfactometer assay. Briefly, 10 μL of a test compound was applied to a 25 × 15 mm filter paper and then placed in one arm of the olfactometer (15 cm base, 10 cm arm length, and 2 cm diameter) as the odorant source. The solvent in the other arm was the mock control. Bee responses within 5 min were scored as “made a choice” when an individual moved at least 2/3 into one arm. More than thirty honeybees were used in each behavioral assay in series of concentration of the test compound.

### Electroantennogram recording

In the EAG assay, both ends of the honeybee antenna were cut and covered with a conductive gel (Parker Laboratories Inc., USA), and the honeybee head was attached to the reference electrode. In a preliminary test, we found that both 9-ODA and HOB are difficult to dissolve in hexane. Thus, we first dissolved them in ethanol and then diluted them with paraffin oil to the desired dose. Ethanol alone, diluted with paraffin oil, was used as a negative control. A 10 μL stimulus was loaded onto a 5.0 × 0.5 cm filter paper strip and then inserted into a syringe with a continuous flow of 500 mL/min and an air humidity of 60-70%. The pulse flow duration was 0.2 s, and the antenna response was recorded for 5 s. To ensure EAG sensitivity restoration, we had 1-minute gaps between two stimulations. For the EAG inhibition test, 9-ODA and HOB were first separated, and then the two pulse flows were mixed at the end of the tube before puffed against the *A. cerana* antennae. The negative control was performed both at the beginning and the end of each preparation. The EAG data were normalized using the negative control data.

### Single sensillum recording

In the SSR test, 12- to 15-day-old drones were wedged into a 1 mL plastic pipette tip, and the protruding head was fixed to the rim of the pipette tip with dental wax. One of the exposed antennae was stuck to a coverslip with double-sided tape under a microscope (LEICA Z16 APO, Germany). The reference tungsten electrode was inserted into the eye, and spikes were recorded by inserting the tungsten electrode into the base of a sensillum until a stable electrical signal with a high signal-to-noise ratio was achieved. For stimulus delivery, 10 μL of the QMP component was added on a 1 cm × 2.5 cm filter paper strip and then inserted into a Pasteur pipette. A flow of purified and humidified air (2 L/min) was continuously maintained on the antennae through a 14-cm-long metal tube controlled (Syntech Hilversum, Netherlands) by a Syntech stimulus controller (CS-55 model, Syntech, Germany). The two antenna were exposed to a stimulus for 500 ms with airflow of 0.6 L/min through a Pasteur pipette. The action potential signals were amplified using a pre-amplifier (IDAC-4 USB System, Syntech, Germany) and visualized by the Autospike 32 software (Syntech, Germany). The number of induced spikes were calculated as the subtraction of the firing spike number by spontaneous spikes number before the stimulus.

### In situ hybridization

Antisense and sense digoxigenin- and biotin-labeled riboprobes of *AcerOr11* were synthesized using linearized pGEMHE plasmids containing appropriate insertion sequences as a template using the DIG and Biotin RNA Labeling Mix (Roche, Germany) and T7 RNA Polymerase (Roche, Germany). Subsequently, the probe was digested into approximately 400 base fragments by incubating in carbonate buffer (80 mM NaHCO_3_, 120 mM Na_2_CO_3_, pH 10.2).

Antennae of 12- to 15-day-old honeybees were collected and then embedded in a Tissue-Tek optimal cutting temperature compound (Sakura Finetek, USA). Longitudinal and transverse sections (10 µm thick) through antennae were prepared using the Cryostar NX50 cryostat (ThermoFisher, USA) at −25 ℃. The sections were thaw-mounted on adhesive microscope slides (Citotest, China) and immediately utilized for in situ hybridization experiments.

Tissues were fixed in 4% paraformaldehyde, and slides were washed with phosphate-buffered saline (PBS) buffer and 0.6% HCl respectively. For pre-hybridization, slides were immersed in 50% formamide with 2× saline-sodium citrate (SSC) for 1 h at 60 ℃. Afterward, the slides were added with 100 μL of the hybridization buffer containing the labeled probe for *AcerOr11* and incubated at 60 ℃ for a minimum of 16 h. After hybridization, slides were washed in 0.2× SSC, followed by treatment with a 1% blocking solution (Roche, Germany) prepared in tris-buffered saline (TBS) buffer with 0.03% Triton X-100. Anti-digoxigenin-AP, Fab fragments, and NBT/BCIP (all from Roche, Germany) were used to detect the DIG-labeled probe under an Upright Microscope BX51 (Olympus, Japan). Streptavidin-Cy3™ (Roche, Germany) was used to detect the biotin-labeled probe, and the fluorescence signals were visualized under a Zeiss LSM 880 confocal microscope (Zeiss, Jena, Germany) using excitation at 550 nm.

### RNA extraction, gene cloning, and quantitative PCR

RNA samples from different tissues, including the antenna, proboscis, thorax, abdomen, and legs, were collected from 15 bees per caste. Total RNA was extracted using the TRIzol reagent (Invitrogen, USA) following the manufacturer’s protocols. The concentration and purity of the extracted RNA were measured by a NanoDrop 2000 spectrophotometer (Thermo Fisher Scientific, USA) and 1% agarose gel electrophoresis, respectively.

For PCRs, we used gene-specific primers for *AcerOrco* and *AcerOr11* (Table S1). First-strand cDNA synthesis was performed using the TransScript One-Step gDNA Removal and cDNA Synthesis SuperMix (Transgen Biotech, Beijing, China). PCR was performed with the TSINGKE TSE101 PCR enzyme mix (TsingKe Biotech, Beijing, China) at the following conditions: 2 min at 98 °C; followed by 35 cycles of 98 °C for 10 s, 50-60 °C for 10 s, and 72 °C for 20 s; and final extension for 5 min at 72 °C. PCR-amplified products were examined and gel purified using the SanPrep Column DNA Gel Extraction Kit (Sangon Bio, Shanghai, China). The purified PCR products were subcloned into a pGEMHE vector between the BamHI and HindIII restriction sites using the pEASY-Uni Seamless Cloning and Assembly Kit (Transgen Biotech, Beijing, China).

For the qPCR assay, the cDNA sample was quantified using 1 μg of total RNA, and *β-actin* was used as an internal control gene. Primers for *AcerOr11* and *AcerOrco* were designed using Primer 3 (https://bioinfo.ut.ee/primer3-0.4.0/) (Table S1). RT-qPCR was conducted on a LightCycler 480 II Detection System (Roche, Switzerland) with TransStar Tip Top Green qPCR Supermix (Transgen Biotech, China) at the following conditions: 94 ℃ for 30 s, followed by 45 cycles of 94 ℃ for 5 s, 55 ℃ for 15 s, and 72 ℃ for 10 s. qPCR data were analyzed by the 2^-ΔΔCT^ method.

### Deorphanization of AcerORs in the *Xenopus* oocyte system

The cRNAs with the templates, the linearized pGEMHE vector containing of *AcerOrco* and *AcerOr11*, using the mMESSAGE mMACHINE T7 Kit (Ambion, USA) following the manufacturer’s instructions. The cRNAs were adjusted concentration of 200ng/μL in nuclease-free water and 18.4 nL of *AcerOr11* with same amount of *AcerOrco* cRNAs were microinjected into *Xenopus laevis* oocytes at vegetal pole in stages V or VI using a NanoLiter 2000 injector (World Precision Instruments, Sarasota, USA). Subsequently, oocytes were incubated at 18 °C for 2-8 days in Barth’s solution (96 mM NaCl, 2 mM KCl, 5 mM MgCl_2_, 0.8 mM CaCl_2,_ and 5 mM HEPES; pH 7.6) supplemented with 50 μg/mL tetracycline, 100 μg/mL streptomycin, and 500 μg/mL sodium pyruvate.

A two-electrode voltage-clamp (TEVC) technique was used to record the ion channel-induced currents in *Xenopus* oocytes at a holding potential of −80 mV. For I-V curves, the holding potentials were held between −80 and +40 mV. Signals were amplified with an Axonclamp 900A amplifier (Molecular Devices, San Jose, USA). Data acquisition and analysis were performed using Axon Digidata 1550B and pCLAMP10 software, using 50 Hz low-pass filters and digitization at 1 kHz (Molecular Devices, USA). The stock solutions (1 M) of all compounds were prepared in DMSO and then diluted with Ringer buffer. Data collected in TEVC was analysis by Clampfit 10 software. *AcerOr11* expressed ORs were deorphanized against a panel of 163 odorants, including honeybee pheromones and plant volatiles (Table S2). The I-V curves were measured by applying a series of voltages, −80, −60, −40, −20, 0, +20, and +40 to the tested eggs and current changes were observed.

## Results

### Queen-released HOB reduces 9-ODA attraction to drones

To measure the countents of QMPs in *A. cerana*, we performed GC-MS analysis on 12- to 15-day-old virgin queens, mated queens as well as drones and workers. 9-ODA and 9-HDA, two main component in QMPs, were detectable in both virgin and mated queens (Figure 1A, B), but interestingly, HOB was only detected in the mated queens other than virgins (Figure 1B). This notice us that HOB might exercise some functions on the post-mating regulation. Notably, the amount of HOB released by mated queens was significantly smaller than that of 9-ODA (Table S3).

**Figure 1.**
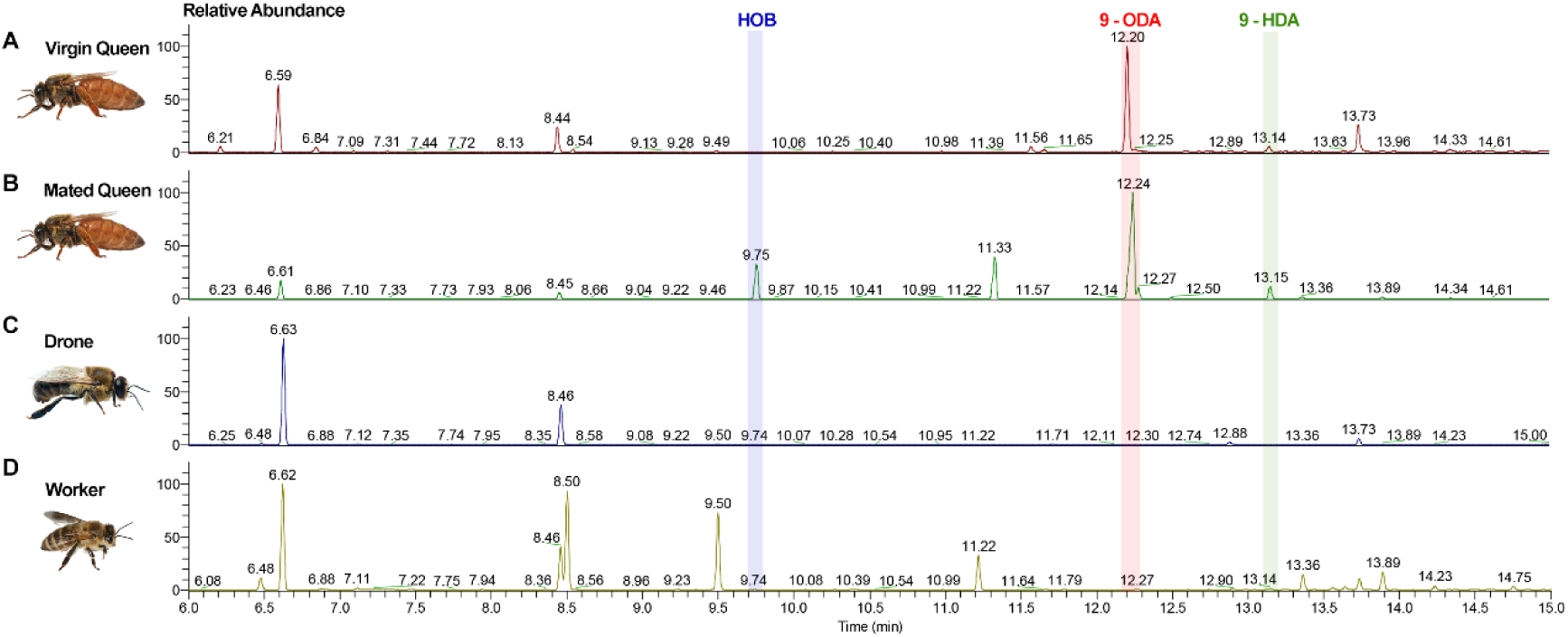
GC-MS analysis of *A. cerana* head extracts. GC-MS analysis of head extracts from (A) 12- to 15-day-old virgin queens, (B) mated queens (mated on day 6 or 7), (C) 12- to 15-day-old drones, and (D) 12- to 15-day-old workers.

Thus we aim to test the behavioral effects of HOB on drones, we used the Y-tube olfactometer assay. First, we used 9-ODA as a stimulus and found that 9-ODA had a significant attractive effect on drones only at the highest concentration (100 μg) were applied (N > 30, *P* < 0.05, two-tailed, T-test). At as lower concentrations as 0.1-10 µg, the attraction effect of 9-ODA was not significant (*P* = 0.0668-0.9580) (Figure 2A). For HOB, it alone did not elicit any effect on drones at all the tested concentrations (0.1-100 μg) (Figure 2B). Then, we mixed 100 µg of 9-ODA with different concentrations of HOB and found 9-ODA’s attraction were suppressed when HOB concentration were beyond 1 µg (*P* = 0.2302-0.9414) (Figure 2C). This suggested that queens only released HOB after mating to compromise the attraction to drones.

**Figure 2.**
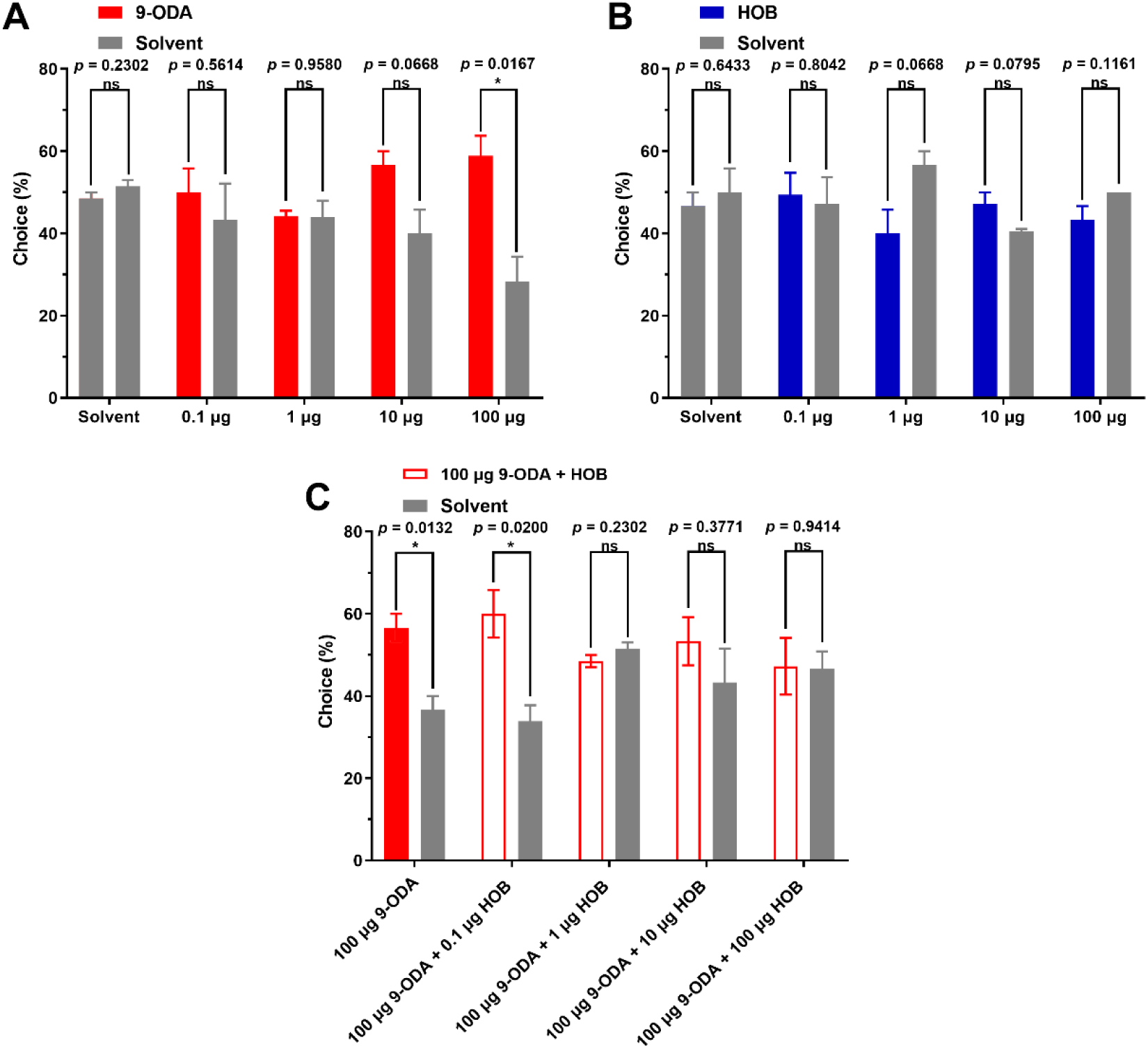
Behavioral responses of *A. cerana* to 9-ODA and HOB. Behavioral responses of drones to (A) 9-ODA alone (N>30, two-tailed, T-test), (B) HOB alone (N>30, two-tailed, T-test), and (C) the mixture (N>30, two-tailed, T-test).

### HOB inhibited 9-ODA neurons in sensilla placodes

To explore the physiological role of HOB *in vivo*, we conducted an EAG assay. First, we tested the olfactory response in the antenna to 9-ODA or HOB alone in different castes. 9-ODA elicited significant EAG responses in drones at the concentration of 100 µg (Figure 3A); the normalized EAG response was 64.63 ± 2.08, which was nearly 64 times higher than that of the negative control group (paraffin oil). The antenna of workers and queens only exhibited a mild response to 9-ODA (N ≥ 5, *P* < 0.05, Wilcoxon signed ranked test) (Figure 3C, S1 A-B). On the contrary, HOB did not elicit any significant antennal response in all castes (Figure 3B and Figure S1).

**Figure 3.**
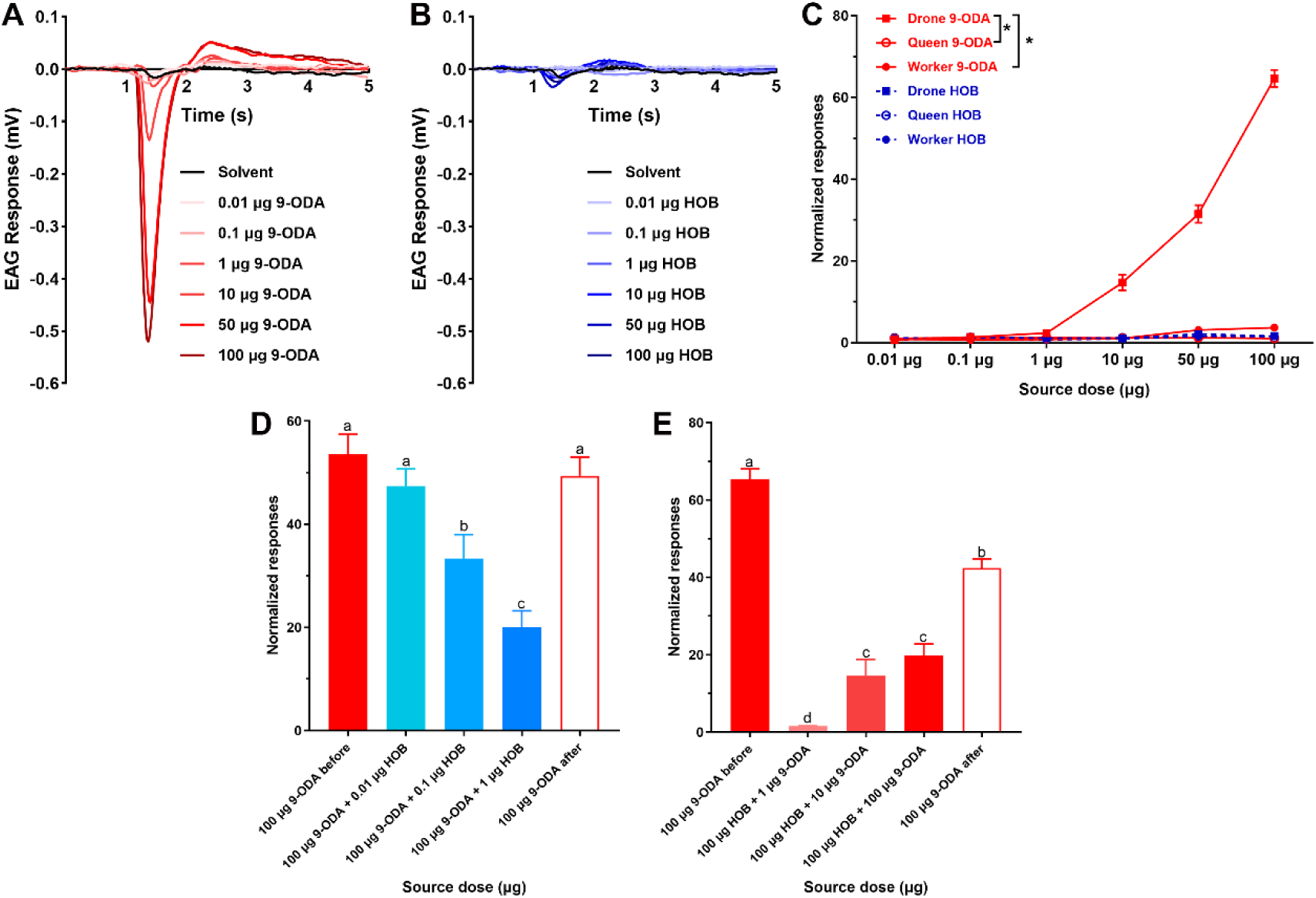
EAG response of 9-ODA and HOB in *A. cerana*. (A, B) EAG responses of drones to increasing doses (0.01 to 100 μg) of 9-ODA and HOB. (C) Dose-response curves for normalized EAG response to 9-ODA and HOB in *A. cerana* (N = 5-9, *P*<0.05, Wilcoxon signed ranked test). (D, E) HOB elicited dose-dependent inhibition of EAG responses to 9-ODA (N = 6-7, *P*<0.05, One-way ANOVA followed by Turkey’s test). Relative EAG responses = Em/CKm, Em represent the mean responses for the test volatile compound and CKm was negative control.

We next stimulated the antenna with 9-ODA-HOB mixtures. The 9-ODA concentration was fixed to 100 µg for a saturated EAG response. We observed an inhibitory effect of HOB on the EAG response to 9-ODA (Figure S2) in a dose-dependent manner. Mixed with 0.1 and 1 µg HOB, the normalized EAG responses to 9-ODA decreased to 33.33±4.34 and 20.05±2.94, respectively (N = 6-7, *P* < 0.05, One-way ANOVA followed by Turkey’s test), which were significantly lower than those from 9-ODA alone (Figure 3D). Intriguingly, HOB at 100 µg eliminated the effect of all tested 9-ODA concentrations (0.01-1 µg; N = 7, *P* < 0.05) (Figure 3E).

To examine the *in vivo* effect of HOB on the ORN response, we conducted single sensillum recordings (SSRs). There are three types of chemosensory sensilla were observed in the *A. cerana* drone’s antenna, including the sensilla trichodea, placodea, and basiconica. Sensilla placodea obviously outnumbers the other two types of sensilla. Of over 150 chemosensory sensilla tested with 9-ODA and HOB, only sensilla placodea showed responses to 9-ODA and HOB, and no response were detected in sensilla trichodea and sensilla basiconica. Consistent with *A. mellifera* studies ^11^, three types of spike amplitudes were observed in *A. cerana* sensilla placodea, suggesting the presence of three ORNs (neurons A, B, and C) (Figure 4A). When stimulated with 9-ODA, the activity of the A neurons was significantly increased compared to the negative control (N = 5, *P* < 0.001, two-tailed, T-test) (Figure 4B, C), when stimulated with HOB, the spontaneous activity of the A neuron was significantly reduced (N = 5, *P* < 0.001), indicating the inverse effect of HOB. Intriguingly, either 9-ODA or HOB changes the spontaneous activity of B and C neuron (Figure 4D), suggesting they were specifically acting on A neuron. We next tested the effect of the 9-ODA-HOB mixture on A neurons. The presence of HOB inhibited 9-ODA-induced activity in A neurons (N = 5, *P* < 0.05) (Figure 4E). Taken together, these results indicate that HOB inhibits not only the antennal response to 9-ODA, but also the spontaneous firing at absence of 9-ODA, implied HOB could act as an inverse agonist at the ORNs level.

**Figure 4.**
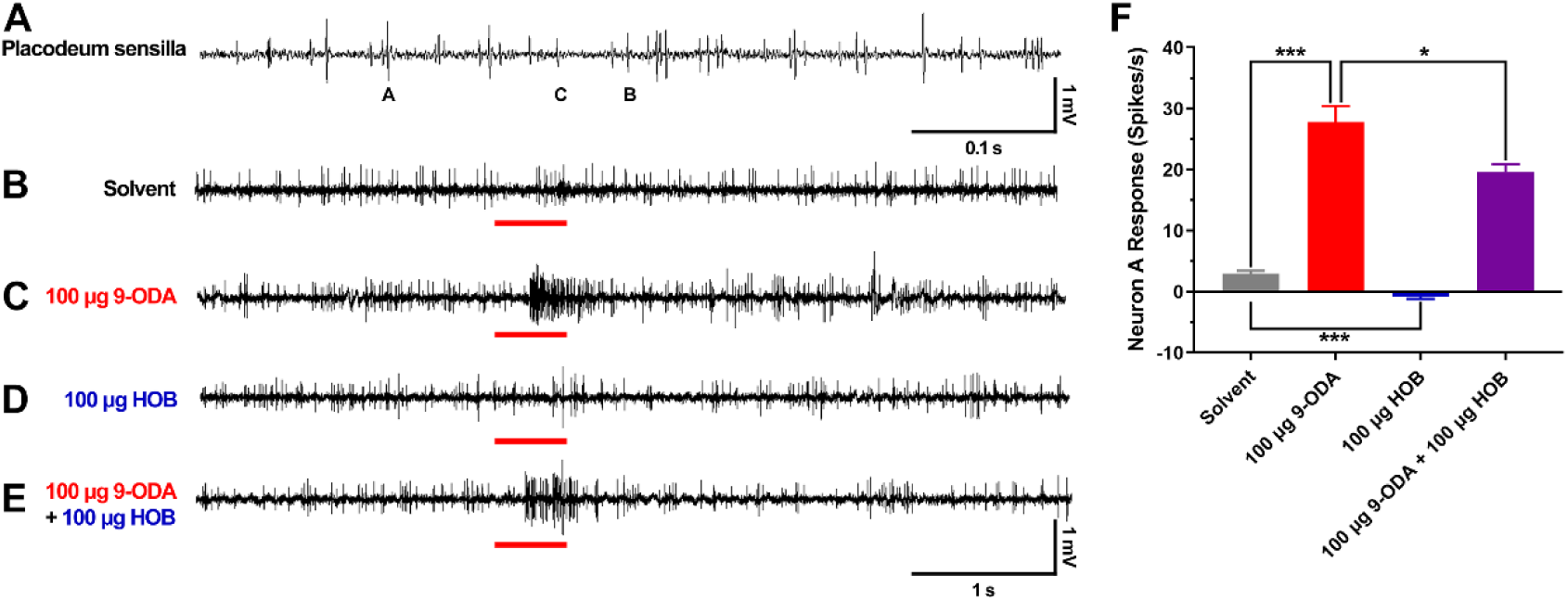
SSR with 9-ODA and HOB in placodeum sensilla. (A-E) Representative traces of placodeum sensilla response to solvent, 100 μg 9-ODA, 100 μg HOB, and the 9-ODA-HOB mixture. (F) Statistic analysis of A neuron’s spikes to 100 μg 9-ODA, 100 μg HOB, and the 9-ODA-HOB mixture (N = 5-6, two-tailed, T-test).

### *AcerOr11* is abundantly expressed in sensilla placodea

To identify the potential molecular mechanism underlying the physiological response to 9-ODA and HOB in the antenna, we conducted an RT-qPCR survey of the *AmelOr11* homolog in *A. cerana* in different tissues and castes. We found that *AcerOr11* is expressed only in the drone antenna, while little expressions were also detected in the queen and worker antennae (Figure S3). This result indicated the *AcerOr11* plays a crucial and special role in the drone’ detecting queen’s mating cue.

For further determining the expression level and location of *AcerOr11*, we conducted *in situ* hybridization experiment. The DIG-labeled riboprobes for *AcerOr11* were applied to transversal antennal sections of bees from the castes. We found that many cells in the drone antenna expressed *AcerOr11*, which was uniformly distributed from the F1 to the F11 segments (Figure 5A, B). Only a few cells in workers expressed *AcerOr11*, while *AcerOr11* expressing were detected in queens (Figure 5C, D, L, M). No labeled cells were observed in the negative control group (Figure S4). *AcerOr11*-labeled area were mainly distributed in dendrite-like structures of ORNs housed in sensilla placodea (Figure 5K). The *AcerOr11* and 4’,6-diamidino-2-phenylindole (DAPI) labeled areas were neatly separated (Figure 5N), suggesting that *AcerOr11* is expressed in the ORN cytoplasm instead of nucleus.

**Figure 5.**
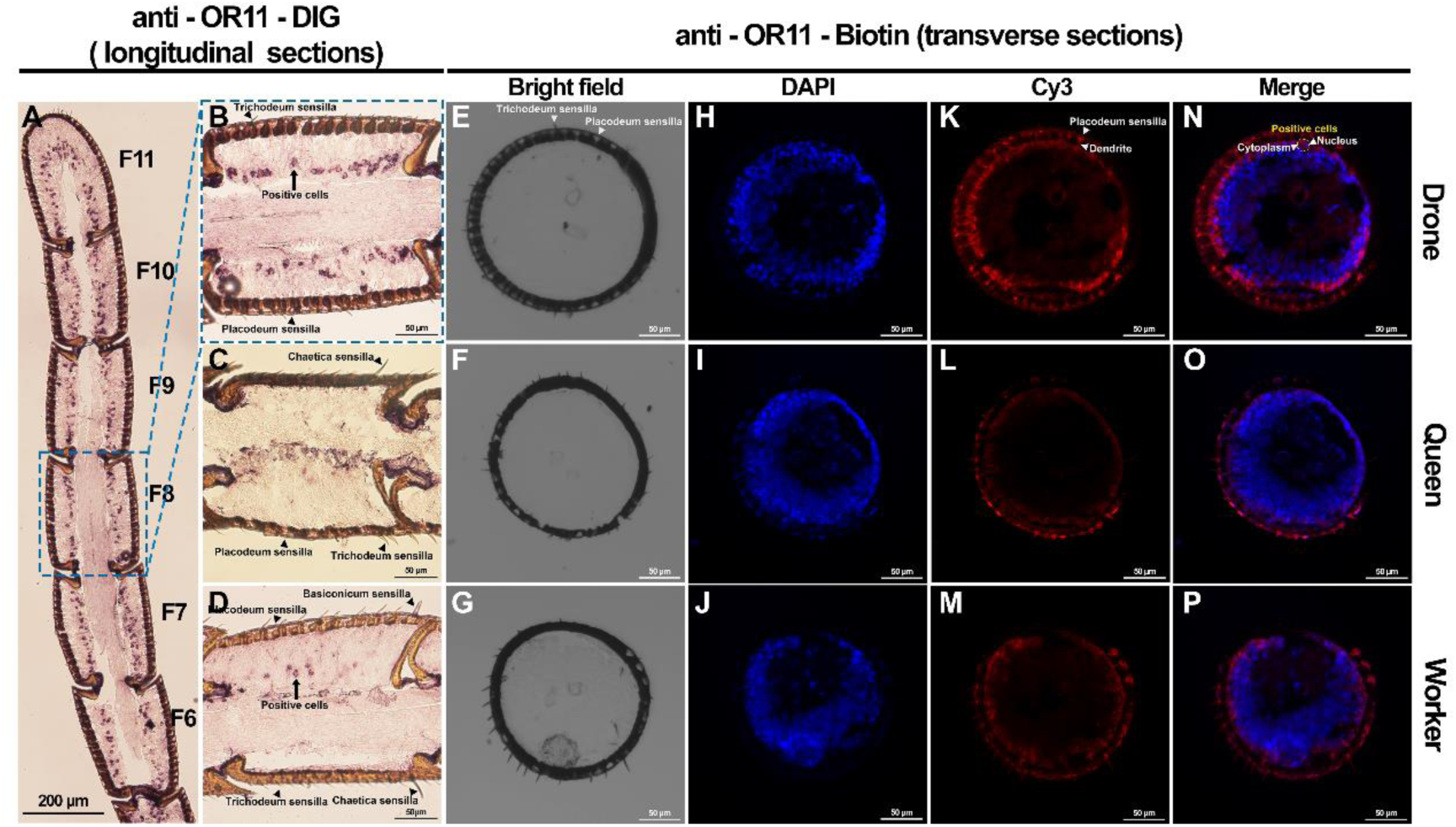
In situ hybridization of AcerOr11. (A-D) Chromogenic in situ hybridization using the anti-Or11-DIG specific probe on longitudinal sections through the median plane of flagellar segments from antennae of (A, B) drones, (C) queens, and (D) workers. (E-P) High magnification images of a two-color FISH experiment on a transverse section incubated with anti-Or11-Biotin for AcerOr11 (K, L, and M; red), and the counterstained images with DAPI (H, I, and J; blue). Bright-field (E, F, and G) and merged (N, O, and P) images are also presented for reference.

### Dual coding of *AcerOr11* against 9-ODA and HOB

To identify the QMP ORs in *A. cerana*, we cloned *AcerOr11*, which is the 1:1 homolog of *AmelOr11* (9-ODA receptor in *A. mellifera*) and specifically expressed in sensilla placodea from drones ^7^. We functionally expressed *AcerOr11* in the *Xenopus laevi*s system and screened them with a 163-compound panel. AcerOr11 produced robust regular currents in response to increasing concentrations of 9-ODA (EC_50_ = 0.35 nM). Meanwhile, HOB elicited inverse currents (Figure S5) in a dose-dependent manner (EC_50_ = 150 nM) (Figure 6A-D).

**Figure 6.**
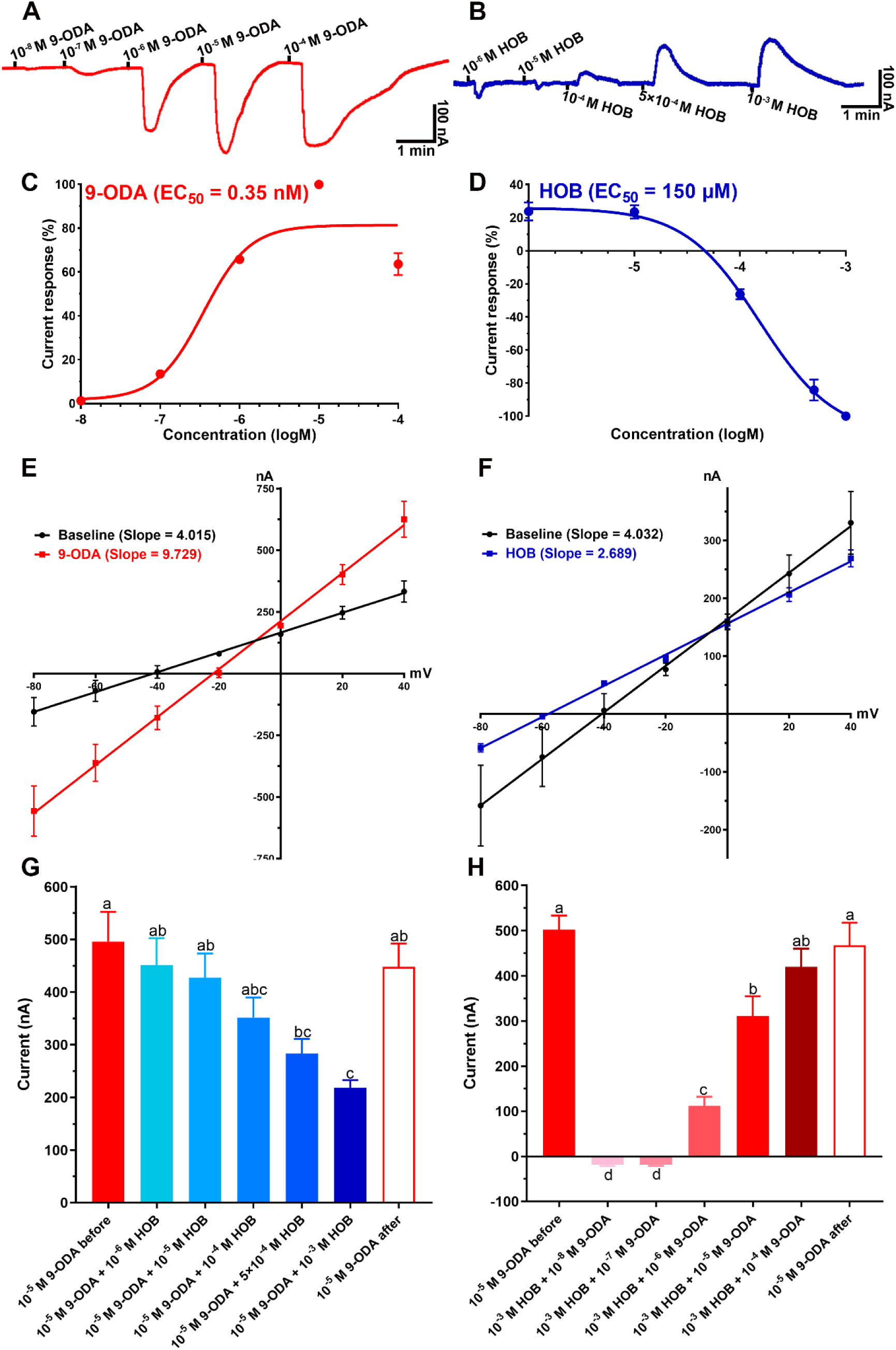
TEVC response of AcerOr11 to 9-ODA and HOB. Representative trace of currents recorded from AcerOr11/AcerOrco expressing oocytes when challenged with increasing doses of (A) 9-ODA from 10^-8^ M to 10^-4^ M and (B) HOB from 10^-6^ M to 10^-3^ M. Dose-response curves for (C) 9-ODA activation of AcerOr11/AcerOrco (EC_50_ = 0.35 nM, mean ± SEM, N = 5-6) and (D) HOB activation of AcerOr11/AcerOrco (EC_50_ = 0.15 μM, mean ± SEM, N = 5-7). (E, F) I-V curves of 9-ODA (red) and HOB (blue) at 10^-5^ M and 10^-3^ M in the voltage range of −80 to +40 mV (N = 6). (G, H) HOB elicited dose-dependent inhibition of 9-ODA-induced responses of AcerOr11/AcerOrco-expressing oocytes (mean ± SEM, N = 5-6, *P*<0.05, One-way ANOVA followed by Turkey’s test).

To further confirm the inverse response of HOB on *AcerOr11*, a current-voltage (I-V) curve was constructed using 9-ODA and HOB as stimuli. The results showed that in the range of −80 to +40 mV, the slope of the 9-ODA-elicited I-V curves was much higher (9.729) than that of the baseline (4.015) (Figure 6E). On the contrary, HOB-generated I-V curves had a much lower slope (2.689) compared with the baseline (4.032) (Figure 6F). These results validated that the presence of the agonist 9-ODA dramatically increases the conductivity of the cell membrane, and channels are massively opened. Contrarily, the presence of HOB decreases the conductivity as much as in the resting state.

We next measured the TEVC response of the 9-ODA-HOB mixture under different dose ratio. We fixed the concentration of 9-ODA to 10^-5^ M for a saturated response with increased the concentration of HOB from 10^-6^ M to 10^-3^ M. We found there was a clear dose-dependent inhibitory effect of HOB on 9-ODA activity (Figure S6). At 5×10^-4^ M or higher, HOB significantly inhibited 9-ODA-elicited currents (N = 5, *P* < 0.05, One-way ANOVA followed by Turkey’s test). At 10^-3^ M HOB concentration, the AcerOr11 response to the 9-ODA-HOB mixture was 218.3 ± 12.9 nA, which was near half of that induced by 9-ODA alone (495.7 ± 50.8 nA). Subsequently, we used 9-ODA alone shortly after the serious of test and found that the response to 9-ODA was recovered (Figure 6G). These results suggest that HOB inhibits the Or11 response to 9-ODA. Likewise, when HOB was at 10^-3^ M and 9-ODA was at lower doses (10^-8^ M and 10^-7^ M), the mixture still induced depolarization (inverse) currents. However, once 9-ODA was raised to 10^-6^ M or higher, the response turned to inverse currents (Figure 6H).

## Discussion

To begin with, GC-MS analysis indicated that HOB was only released by mated queens, which is consistent with a report about HOB in *A. mellifera* queens. In our case, HOB alone did not elicit any behavioral effect on drones, even at the highest doses, in the binary choice assay. However, it significantly reduced the attraction of drones to 9-ODA. Therefore, HOB takes an inverse behavioral effect compared with 9-ODA in *A. cerana*. Previous study showed the antennal-specific protein 1 (*Asp1*), has a high affinity for HOB, and the *Asp1* expression level is positively correlated with colony sizes in both *A. cerana* and *A. mellifera* ^12^, these results support that HOB alone could be detected by *Apis* honeybees at a peripheral olfactory sensing level, and might played crucial role in regulating population dynamics in honeybees. Our behavioral assay results indicate that HOB directly inhibits the mating behavior of drones and might prevents the bee colony from having an oversized population, trading off the number of offspring for the allocation of energy resources, including physical resources and parental care.

In insects, excitatory and inverse chemical signals are equally important for maintaining the population dynamic. In *Helicoverpa armigera*, female moths release cis-11-Hexadecenol (Z11-16:OH) to repel males and avoid non-optimal mating ^13^. Likewise, in *Drosophila*, Gr8-associated alkenes inhibit courtship behaviors ^14^. Collectively, these findings suggest that the inverse agonist may be widespread in insects. With a few exceptions, honeybee queens do not remate in her lifetime ^15^. Mated queens release HOB to prevent drones from multiple mating, which can help in avoiding over-sized populations and resource waste due to multiple mating and subsequent breeding.

Interreceptor inhibition by semiochemicals at the sensillum level, a well-established phenomenon in *D. melanogaster*, is mediated by non-synaptic “lateral inhibitions” between neurons located in the same sensilla, termed ephaptic coupling ^16,17^. In mosquitoes, high concentrations of ammonia elicit atypical bursts of action potentials, followed by inhibition in multiple adjacent ORNs ^18^. In *C. quinquefasciatus*, eucalyptol significantly decreases the number of spikes in the ab7 sensillum ^19^. Besides the interreceptor inhibition, here, we show evidence that HOB reduces the activity of adjacent 9-ODA-activated at same receptor of ORNs, supporting the hypothesis of the intrareceptor inhibition. Notably, HOB alone did not elicit any “normal” EAG response, which negates the possibility of any HOB-activated ORs, i.e., HOB does not act as an agonist. Since HOB reduces spontaneous spikes, it had an opposite effect to 9-ODA, that induced the firing of ORNs, we could speculate that HOB act as an inverse agonist to modulate mating behavior at ORNs level.

The *A. cerana AcerOr11* gene is expressed in neurons specifically tuned to 9-ODA and HOB. However, how those two opposite ligands affect the *AcerOr11* receptor is unknown. Therefore, we cloned and expressed *AcerOR11* in the *Xenopus* oocyte system, which exhibited robust responses to 9-ODA while HOB evoked a reverse, concentration-dependent current. These results again indicated that HOB acts as an inverse agonist of *AcerOr11*. Inverse agonists have been identified in various receptor-ligand interactions, including GABAA, melanocortin receptors, mu-opioid receptors, adrenoceptors, and histamine receptors ^20–23^. Currently, the concept of inverse agonists in insect ORs is poorly understood and much less studied. Some experimental evidence suggests that insect ORs have specific inverse agonists. For example, in *C. quinquefasciatus*, OR32 produces regular currents when stimulated with methyl salicylate, while eucalyptol elicits inverse currents, suggesting that eucalyptol might be an inverse agonist ^19^. In *Aedes aegypti*, *AaegOR8* was found to be sensitively tuned to (*R*)-1-octen-3-ol while the structurally unrelated odorant indole inhibited octenol-activated *OR8* and repelled mosquitoes ^24^. The potential mechanism of inverse currents can be speculated in three stages: when the receptor is challenged, the agonist interacts with the ligand binding site, triggering an inward current, while the antagonist blocks the binding site to prevent the agonist binding while maintaining spontaneous activity, and inverse agonist “snare” receptor to inactiveness. Our results suggest that HOB to *AcerOR11* generates an additional signal coding, which may execute more complex regulation instead of a simple “on-off” function.

Agonist-inverse agonist combinations may elicit opposite physiological effects. For example, the mushroom psychoactive compound muscimol induces a relaxing effect by activating the GABAA receptor. On the other hand, beta-carbolines act as inverse agonists and cause convulsive or anxiogenic effects ^20^. Similarly, 9-ODA promotes mating behavior, while HOB inhibits mating. In summary, our results suggest that mating in honeybees is modulated by the two key QMP components, 9-ODA and HOB. *Or11* is not merely an on/off switch but rather functions as a molecular olfactory dimmer.

*A. cerana* is an excellent pollinator for a wide range of host plants and thereby plays an irreplaceable role in the ecosystem and agriculture. Unfortunately, the massive use of pesticides and the introduction of *A. mellifera* in China have reduced its colonies by 80% ^25^. The profound effect of HOB on *A. cerana* mating behavior can be used to regulate apiculture and the conservation of this species.

## Supporting information

Supplemental Files for manuscirpt

## Acknowledgments

We thank Dr. Pingxi Xu (Department of MCB, UC Davis) for critically reading an earlier manuscript draft; Dr. Yuanhong Wang (School of Chemistry, Northeast Normal University) for the help with GC-MS analysis; Xiuli Wang and Fan Zhang (School of life sciences, Northeast Normal Univerisity) for assistance in in situ hybridization; Dr. Baiwei Ma (Chinese Academy of Agricultural Sciences), Dr. Shuai Liu (Department of Plant Protection, Jilin Agricultural University) and Cuiwei Liu for the guidance in SSR experiments.

## Author contributions

Y. W. and J. D. B. designed this study; H. K., S. D., and X. M. performed this research; C.X. provided and reared the honeybee colony; Y. W. and H. K. analyzed the data; Y. W., J. D. B, and B. R. wrote and revised manuscript.

## Declaration of competing interest

All authors declare no conflicts of interest.

## Ethics approval and consent to participate

Not applicable.

## Consent for publication

All authors give their consent to publish this study.

## Availability of data and material

All data used during this study are included in this article and Supplementary materials.

## Funding

This work was supported by the National Key Research and Development Program “Intergovernmental international cooperation on science, technology and innovation” (No. 2023YFE0113600), National Natural Science Foundation (No. 31972289), Israel Science Foundation grants (No. 719/21) and Natural Science Foundation of Jilin Province (No. 20200402001NC and No. 20190301047NY).

## Notes

### Competing Interest Statement

The authors have declared no competing interest.

## References

1 Eckert, C. D., Winston, M. L. & Ydenberg, R. C. The Relationship between Population Size, Amount of Brood, and Individual Foraging Behavior in the Honey Bee, *Apis Mellifera* L. Oecologia 97, 248–255, doi:Doi 10.1007/Bf00323157 (1994).

2 Khoury, D. S., Myerscough, M. R. & Barron, A. B. A quantitative model of honey bee colony population dynamics. PLoS One 6, e18491, doi:10.1371/journal.pone.0018491 (2011).

3 Martin, S., Chaline, N., Drijfhout, F. & Jones, G. Role of esters in egg removal behaviour in honeybee (*Apis mellifera*) colonies. Behav Ecol Sociobiol 59, 24–29, doi:10.1007/s00265-005-0004-0 (2005).

4 Le Conte, Y., Mohammedi, A. & Robinson, G. E. Primer effects of a brood pheromone on honeybee behavioural development. Proc Biol Sci 268, 163–168, doi:10.1098/rspb.2000.1345 (2001).

5 Sandoz, J. C., Deisig, N., de Brito Sanchez, M. G. & Giurfa, M. Understanding the logics of pheromone processing in the honeybee brain: from labeled-lines to across-fiber patterns. Front Behav Neurosci 1, 5, doi:10.3389/neuro.08.005.2007 (2007).

6 Plettner, E. et al. Species- and caste-determined mandibular gland signals in honeybees (*Apis*). J Chem Ecol 23, 363–377, doi:DOI 10.1023/B:JOEC.0000006365.20996.a2 (1997).

7 Wanner, K. W. et al. A honey bee odorant receptor for the queen substance 9-oxo-2-decenoic acid. Proc Natl Acad Sci U S A 104, 14383–14388, doi:10.1073/pnas.0705459104 (2007).

8 Sandoz, J. C. Odour-evoked responses to queen pheromone components and to plant odours using optical imaging in the antennal lobe of the honey bee drone *Apis mellifera* L. J Exp Biol 209, 3587–3598, doi:10.1242/jeb.02423 (2006).

9 McKenzie, S. K., Fetter-Pruneda, I., Ruta, V. & Kronauer, D. J. Transcriptomics and neuroanatomy of the clonal raider ant implicate an expanded clade of odorant receptors in chemical communication. Proc Natl Acad Sci U S A 113, 14091–14096, doi:10.1073/pnas.1610800113 (2016).

10 Song, X. M. et al. Various Bee Pheromones Binding Affinity, Exclusive Chemosensillar Localization, and Key Amino Acid Sites Reveal the Distinctive Characteristics of Odorant-Binding Protein 11 in the Eastern Honey Bee, Apis cerana. Front Physiol 9, 422, doi:ARTN 422 10.3389/fphys.2018.00422 (2018).

11 Kaissling, K. E. & Renner, M. Antennale Rezeptoren für Queen Substance und Sterzelduft bei der Honigbiene. Ztschrift Für Verglchende Physiologie 59, 357–361 (1968).

12 Wu, F. et al. Differences in ASP1 expression and binding dynamics to queen mandibular pheromone HOB between *Apis mellifera* and *Apis cerana* workers reveal olfactory adaptation to colony organization. Int J Biol Macromol 217, 583–591, doi:10.1016/j.ijbiomac.2022.07.064 (2022).

13 Chang, H. et al. A Pheromone Antagonist Regulates Optimal Mating Time in the Moth *Helicoverpa armigera*. Curr Biol 27, 1610–1615, doi:10.1016/j.cub.2017.04.035 (2017).

14 Vernier, C. L. et al. A pleiotropic chemoreceptor facilitates the production and perception of mating pheromones. iScience 26, 105882, doi:10.1016/j.isci.2022.105882 (2023).

15 Boomsma, J. J., Baer, B. & Heinze, J. The evolution of male traits in social insects. Annu Rev Entomol 50, 395–420, doi:10.1146/annurev.ento.50.071803.130416 (2005).

16 Su, C. Y., Menuz, K., Reisert, J. & Carlson, J. R. Non-synaptic inhibition between grouped neurons in an olfactory circuit. Nature 492, 66–72, doi:10.1038/nature11712 (2012).

17 Zhang, Y. et al. Asymmetric ephaptic inhibition between compartmentalized olfactory receptor neurons. Nat Commun 10, 1560, doi:ARTN 1560 10.1038/s41467-019-09346-z (2019).

18 Clark, J. T. et al. Chemosensory detection of aversive concentrations of ammonia and basic volatile amines in insects. iScience 26, 105777, doi:10.1016/j.isci.2022.105777 (2023).

19 Xu, P. et al. Odorant Inhibition in Mosquito Olfaction. iScience 19, 25–38, doi:10.1016/j.isci.2019.07.008 (2019).

20 Sieghart, W. Pharmacology of benzodiazepine receptors: an update. J Psychiatry Neurosci 19, 24–29 (1994).

21 Ollmann, M. M., Lamoreux, M. L., Wilson, B. D. & Barsh, G. S. Interaction of Agouti protein with the melanocortin 1 receptor in vitro and in vivo. Genes Dev 12, 316–330, doi:10.1101/gad.12.3.316 (1998).

22 Wang, D. X., Raehal, K. M., Bilsky, E. J. & Sadee, W. Inverse agonists and neutral antagonists at mu opioid receptor (MOR): possible role of basal receptor signaling in narcotic dependence. J Neurochem 77, 1590–1600, doi:DOI 10.1046/j.1471-4159.2001.00362.x (2001).

23 Khilnani, G. & Khilnani, A. K. Inverse agonism and its therapeutic significance. Indian J Pharmacol 43, 492–501, doi:10.4103/0253-7613.84947 (2011).

24 Dekel, A., Sar-Shalom, E., Vainer, Y., Yakir, E. & Bohbot, J. D. The ovipositor cue indole inhibits animal host attraction in *Aedes aegypti* (Diptera: Culicidae) mosquitoes. Parasit Vectors 15, 422, doi:10.1186/s13071-022-05545-8 (2022).

25 Yang, G. Protect and develop *Apis cerana cerana* to maintain ecological balance. Apiculture of China 70, 18–20 (2019).

