## Supplemental Files for manuscirpt for "The dual coding of a single sex pheromone receptor in regulating the mating behavior of Asian Honeybee *Apis cerana*"

Haoqin Ke *et al.*

**This PDF file includes:**

Table S1 to S3
Figures S1 to S6

**Table S1. Primers list in this study.**

| Purpose | Primer | Primer (5' - 3') |
|---|---|---|
| Gene cloning | Orco-F | ACGTACCCGCCTCAA |
| | Orco-R | TCATTTAGTCACAGTTTCCA |
| | Or11-F | TCACGAACAAGCTTTCATCGG |
| | Or11-R | GAAAGTGAACAAAGTGCTGTGTACA |
| Infusion cloning | Orco-F | AATTCCCCGGGGATCCACCATGATGAAGTTCAAGCAACAGGG |
| | Orco-R | TAACCAGATCAAGCTTTCACTTCAGTTGCACCAACACC |
| | Or11-F | AATTCCCCGGGGATCCACCATGGTCCAAATTAGAAACGCGAAAG |
| | Or11-R | TAACCAGATCAAGCTTTTACGTAACTGTACGTAACATATTC |
| qPCR | Orco-F | GGATCAGAGGAGGCCAAAAC |
| | Orco-R | CCAACACCGAAGCAAAGAGA |
| | Or11-F | ATGTGCGGTTTGCTGAAGA |
| | Or11-R | CGAGAAGGTGCCAATGACG |
| | *β-actin*-F | GGCTCCCGAAGAACATCC |
| | *β-actin*-R | TGCGAAACACCGTCACCC |
| In situ hybridization | Or11-F | AATTCCCCGGGGATCCACCTTACGTAACTGTACGTAACATATTC |
| | Or11-R | TAACCAGATCAAGCTTATGGTCCAAATTAGAAACGCGAAAG |

Note: The underlined DNA sequence was the enzyme restriction site.

**Table S2. Compounds used for deorphanize the *AcerOr11*.**

| Compound | Cas No. | Source | Company | Country | Purity | Structure |
|---|---|---|---|---|---|---|
| (*E*)-9-Oxo-2-decenoic acid (9-ODA) | 334-20-3 | Queen retinue pheromone | Chem-strong Biotechnology Co., LTD | Shenzhen, China | 95.0 % | 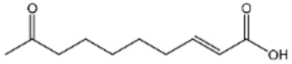 |
| 9-Hydroxy-2-decenoic acid (9-HDA) | 4448-33-3 | Queen retinue pheromone | Albs Biotechnology Co., LTD | Changsha, China | 95.0 % | 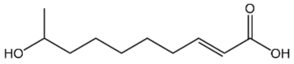 |
| Methyl *p*-hydroxybenzoate (HOB) | 99-76-3 | Queen retinue pheromone | Rhawn Biotechnology Co., LTD | Shanghai, China | 99.0 % | 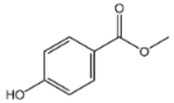 |
| 4-Hydroxy-3-methoxyphenylethanol (HVA) | 2380-78-1 | Queen retinue pheromone | Rhawn Biotechnology Co., LTD | Shanghai, China | 98.0 % | 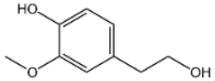 |
| (E)-10-Hydroxy-dec-2-enoic acid (10-HDA) | 765-01-5 | Queen retinue pheromone | Rhawn Biotechnology Co., LTD | Shanghai, China | 98.0 % | 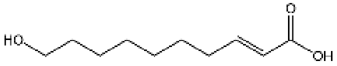 |
| 10-Hydroxy-decanoic acid (10-HDAA) | 1679-53-4 | Queen retinue pheromone | Rhawn Biotechnology Co., LTD | Shanghai, China | 96.0 % | 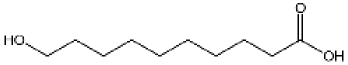 |
| Methyl oleate (MO) | 112-62-9 | Queen retinue pheromone | Aladdin Biochemical Technology Co., LTD | Shanghai, China | 99.0 % | 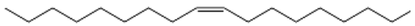 |
| 1-Hexadecanol (PA) | 36653-82-4 | Queen retinue pheromone | Aladdin Biochemical Technology Co., LTD | Shanghai, China | 99.0 % | 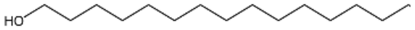 |
| Linolenic acid (LA) | 463-40-1 | Queen retinue pheromone | Merda Technology Co., LTD | Beijing, China | 99.0 % | 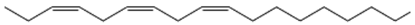 |
| Geraniol | 106-24-1 | Nasonov pheromone | Sigma-Aldrich LLC. | MO, USA | 98.0 % | 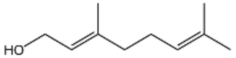 |
| Geranial | 5392-40-5 | Nasonov pheromone | Aladdin Biochemical Technology Co., LTD | Shanghai, China | 97.0 % | 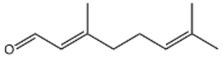 |

| **Compound** | **Cas No.** | **Source** | **Company** | **Country** | **Purity** | **Structure** |
|---|---|---|---|---|---|---|
| Farnesol | 4602-84-0 | Nasonov pheromone | Bidepharm Medical Technology Co., LTD | Shanghai, China | 95.0% | 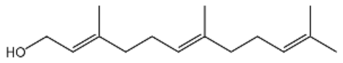 |
| Nerol | 106-25-2 | Nasonov pheromone | Rhawn Biotechnology Co., LTD | Shanghai, China | 97.0% | 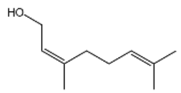 |
| Geranic acid | 459-80-3 | Nasonov pheromone | Rhawn Biotechnology Co., LTD | Shanghai, China | 90.0% | 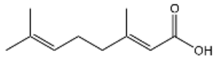 |
| (E)-Citral | 141-27-5 | Nasonov pheromone | Kaiwei Biotechnology Co., LTD | Liaoning, China | 95.0% | 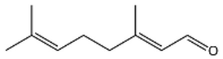 |
| Benzyl acetate | 140-11-4 | Alarm pheromone | Rhawn Biotechnology Co., LTD | Shanghai, China | 99.0% | 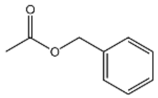 |
| Isoamyl acetate | 123-92-2 | Alarm pheromone | Aladdin Biochemical Technology Co., LTD | Shanghai, China | 99.0% | 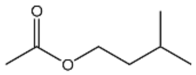 |
| Butyl acetate | 123-86-4 | Alarm pheromone | Aladdin Biochemical Technology Co., LTD | Shanghai, China | 99.7% | 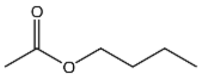 |
| Hexyl acetate | 142-92-7 | Alarm pheromone | Sigma-Aldrich LLC. | MO, USA | 99.0% | 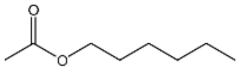 |
| Octyl acetate | 112-14-1 | Alarm pheromone | Rhawn Biotechnology Co., LTD | Shanghai, China | 98.0% | 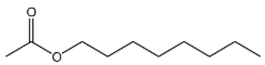 |
| Decyl acetate | 112-17-4 | Alarm pheromone | Aladdin Biochemical Technology Co., LTD | Shanghai, China | 98.0% | 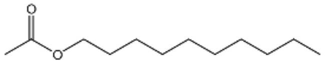 |
| Phenyl acetate | 122-79-2 | Alarm pheromone | Aladdin Biochemical Technology Co., LTD | Shanghai, China | 98.0% | 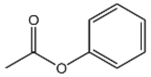 |

| Compound | Cas No. | Source | Company | Country | Purity | Structure |
|---|---|---|---|---|---|---|
| Octanoic acid | 124-07-2 | Alarm pheromone | Aladdin Biochemical Technology Co., LTD | Shanghai, China | 98.0 % | 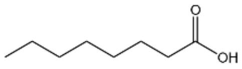 |
| Benzoic acid | 65-85-0 | Alarm pheromone | Aladdin Biochemical Technology Co., LTD | Shanghai, China | 99.5 % | 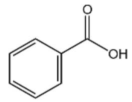 |
| Phenol | 108-95-2 | Alarm pheromone | Aladdin Biochemical Technology Co., LTD | Shanghai, China | 99.0 % | 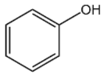 |
| *p*-Cresol | 106-44-5 | Alarm pheromone | Aladdin Biochemical Technology Co., LTD | Shanghai, China | 99.0 % | 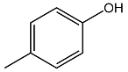 |
| 1-Hexanol | 111-27-3 | Alarm pheromone | Aladdin Biochemical Technology Co., LTD | Shanghai, China | 95.0 % | 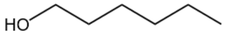 |
| 1-Butanol | 71-36-3 | Alarm pheromone | Rhawn Biotechnology Co., LTD | Shanghai, China | 99.0 % | 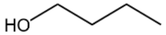 |
| 1-Pentanol | 71-41-0 | Alarm pheromone | Aladdin Biochemical Technology Co., LTD | Shanghai, China | 99.5 % | 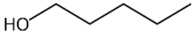 |
| 3-Methyl-1-butanol | 123-51-3 | Alarm pheromone | Rhawn Biotechnology Co., LTD | Shanghai, China | 98.0 % | 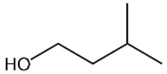 |
| 1-Octanol | 111-87-5 | Alarm pheromone | Aladdin Biochemical Technology Co., LTD | Shanghai, China | 99.5 % | 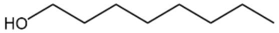 |
| 1-Octadecanol | 112-92-5 | Alarm pheromone | Rhawn Biotechnology Co., LTD | Shanghai, China | 99.0 % | 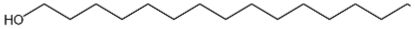 |
| 1-Eicosanol | 629-96-9 | Alarm pheromone | Bidepharm Medical Technology Co., LTD | Shanghai, China | 98.0 % | 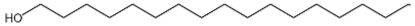 |

| Compound | Cas No. | Source | Company | Country | Purity | Structure |
|---|---|---|---|---|---|---|
| 2-Heptanone | 110-43-0 | Alarm pheromone | Sigma-Aldrich LLC. | MO, USA | 99.0% | 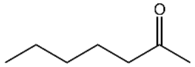 |
| trans-2-Octen-1-ol | 18409-17-1 | Alarm pheromone | Sigma-Aldrich LLC. | MO, USA | 97.0% | 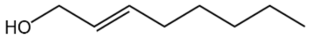 |
| 2-Nonanol | 628-99-9 | Alarm pheromone | Rhawn Biotechnology Co., LTD | Shanghai, China | 98.0% | 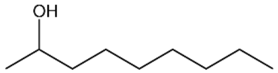 |
| 2-Aminoacetophenone | 551-93-9 | Queen feces pheromone | Bidepharm Medical Technology Co., LTD | Shanghai, China | 99.5% | 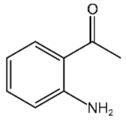 |
| Decyl decanoate | 1654-86-0 | Tergum pheromone | Bidepharm Medical Technology Co., LTD | Shanghai, China | 98.0% | 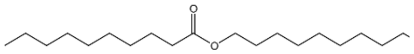 |
| Methyl stearate | 112-61-8 | Brood pheromone | Aladdin Biochemical Technology Co., LTD | Shanghai, China | 99.0% | 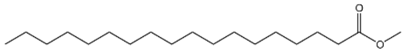 |
| Ethyl stearate | 111-61-5 | Brood pheromone | Macklin Biochemical Technology Co., LTD | Shanghai, China | 99.0% | 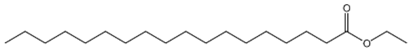 |
| Methyl palmitate | 112-39-0 | Brood pheromone | Aladdin Biochemical Technology Co., LTD | Shanghai, China | 99.0% | 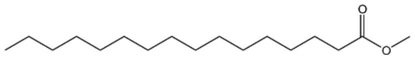 |
| Ethyl palmitate | 628-97-7 | Brood pheromone | Aladdin Biochemical Technology Co., LTD | Shanghai, China | 98.0% | 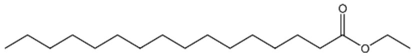 |
| Ethyl oleate | 111-62-6 | Brood pheromone | BioDuly Biotechnology LTD | Nanjing, China | 98.0% | 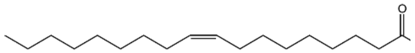 |
| Methyl linoleate | 112-63-0 | Brood pheromone | Bidepharm Medical Technology Co., LTD | Shanghai, China | 100.0% | 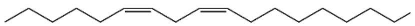 |
| Ethyl linoleate | 544-35-4 | Brood pheromone | Rhawn Biotechnology Co., LTD | Shanghai, China | 97.0% | 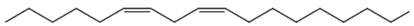 |

| Compound | Cas No. | Source | Company | Country | Purity | Structure |
|---|---|---|---|---|---|---|
| *β*-Ocimene | 13877-91-3 | Brood pheromone | Sigma-Aldrich LLC. | MO, USA | 90.0 % | 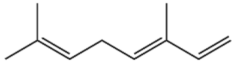 |
| Myrcene | 123-35-3 | Plant volatiles | Rhawn Biotechnology Co., LTD | Shanghai, China | 90.0 % | 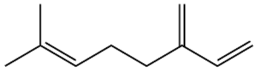 |
| 3-Carene | 13466-78-9 | Plant volatiles | TCI Development Co., LTD | Shanghai, China | 90.0 % | 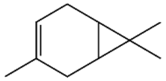 |
| DL-Menthol | 1490-04-6 | Plant volatiles | Aladdin Biochemical Technology Co., LTD | Shanghai, China | 98.0 % | 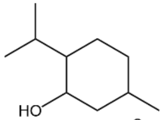 |
| *β*-Ionone | 14901-07-6 | Plant volatiles | Sigma-Aldrich LLC. | MO, USA | 96.0 % | 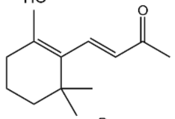 |
| Methyl salicylate | 119-36-8 | Plant volatiles | JK scientific | Beijing, China | 99.0 % | 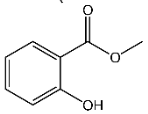 |
| 3,4-Dimethylbenzaldehyde | 5973-71-7 | Plant volatiles | Aladdin Biochemical Technology Co., LTD | Shanghai, China | 97.0 % | 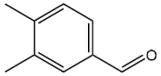 |
| *β*-Citronellol | 106-22-9 | Plant volatiles | Sigma-Aldrich LLC. | MO, USA | 95.0 % | 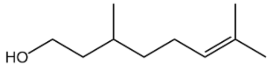 |
| 1,8-Cineole | 470-82-6 | Plant volatiles | Aladdin Biochemical Technology Co., LTD | Shanghai, China | 99.5 % | 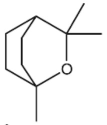 |
| Phenylacetaldehyde | 122-78-1 | Plant volatiles | Sigma-Aldrich LLC. | MO, USA | 90.0 % | 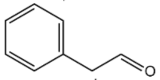 |
| 2-Isobutyl-3-methoxypyrazine | 24683-00-9 | Plant volatiles | Aladdin Biochemical Technology Co., LTD | Shanghai, China | 99.0 % | 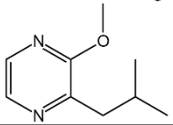 |

| Compound | Cas No. | Source | Company | Country | Purity | Structure |
|---|---|---|---|---|---|---|
| Dodecanoic acid | 143-07-7 | Plant volatiles | Aladdin Biochemical Technology Co., LTD | Shanghai, China | 98.0 % | 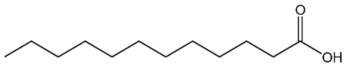 |
| Tetradecanoic acid | 544-63-8 | Plant volatiles | Aladdin Biochemical Technology Co., LTD | Shanghai, China | 98.0 % | 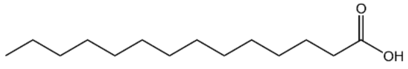 |
| Hexadecanoic acid | 57-10-3 | Plant volatiles | Aladdin Biochemical Technology Co., LTD | Shanghai, China | 98.0 % | 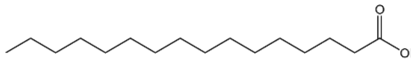 |
| Oleic acid | 112-80-1 | Plant volatiles | Aladdin Biochemical Technology Co., LTD | Shanghai, China | 98.0 % | 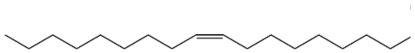 |
| Linoleic acid | 60-33-3 | Plant volatiles | Aladdin Biochemical Technology Co., LTD | Shanghai, China | 95.0 % | 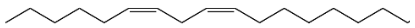 |
| Stearic acid | 57-11-4 | Plant volatiles | Acmec Biochemical Technology Co., LTD | Shanghai, China | 98.0 % | 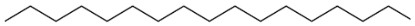 |
| Methyl myristate | 124-10-7 | Plant volatiles | Aladdin Biochemical Technology Co., LTD | Shanghai, China | 98.0 % | 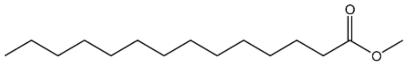 |
| Dodecanal | 112-54-9 | Plant volatiles | Aladdin Biochemical Technology Co., LTD | Shanghai, China | 95.0 % | 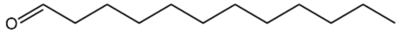 |
| Octadecanal | 638-66-4 | Plant volatiles | Aladdin Biochemical Technology Co., LTD | Shanghai, China | 97.0 % | 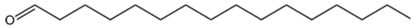 |
| Oleamide | 301-02-0 | Plant volatiles | Bidepharm Medical Technology Co., LTD | Shanghai, China | 98.0 % | 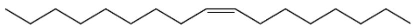 |
| Linalool | 78-70-6 | Plant volatiles | Sigma-Aldrich LLC. | MO, USA | 97.0 % | 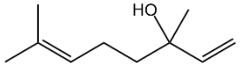 |

| **Compound** | **Cas No.** | **Source** | **Company** | **Country** | **Purity** | **Structure** |
|---|---|---|---|---|---|---|
| 3,7-Dimethyl-1-octanol | 106-21-8 | Plant volatiles | Bidepharm Medical Technology Co., LTD | Shanghai, China | 98.0 % | 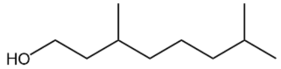  |
| Citronellal | 106-23-0 | Plant volatiles | Heowns Chemical TecHhnology Co., LTD | Wuhan, China | 96.0 % | 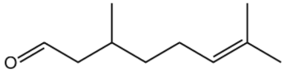  |
| Eugenol | 97-53-0 | Plant volatiles | Sigma-Aldrich LLC. | MO, USA | 99.0 % | 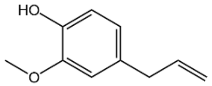  |
| Geranyl acetate | 105-87-3 | Plant volatiles | Bidepharm Medical Technology Co., LTD | Shanghai, China | 95.0 % | 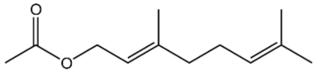  |
| *α*-Methylcinnamaldehyde | 101-39-3 | Plant volatiles | Aladdin Biochemical Technology Co., LTD | Shanghai, China | 95.0 % | 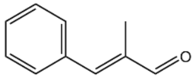  |
| Benzaldehyde | 100-52-7 | Plant volatiles | Aladdin Biochemical Technology Co., LTD | Shanghai, China | 99.5 % | 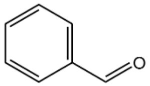  |
| Octanal | 124-13-0 | Plant volatiles | Sigma-Aldrich LLC. | MO, USA | 99.0 % | 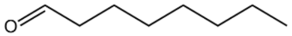  |
| Cis-jasmone | 488-10-8 | Plant volatiles | Aladdin Biochemical Technology Co., LTD | Shanghai, China | 98.0 % | 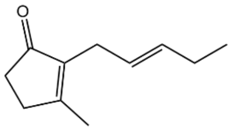  |
| Indole | 120-72-9 | Plant volatiles | Aladdin Biochemical Technology Co., LTD | Shanghai, China | 99.0 % | 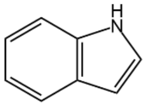  |
| 1- Nonanol | 143-08-8 | Plant volatiles | Bidepharm Medical Technology Co., LTD | Shanghai, China | 98.0 % | 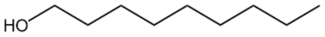  |
| 1-Octen-3-ol | 3391-86-4 | Plant volatiles | Aladdin Biochemical Technology Co., LTD | Shanghai, China | 98.0 % | 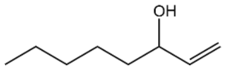  |

| Compound | Cas No. | Source | Company | Country | Purity | Structure |
|---|---|---|---|---|---|---|
| β-Caryophllene | 87-44-5 | Plant volatiles | Bidepharm Medical Technology Co., LTD | Shanghai, China | 90.0 % | 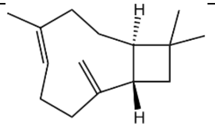 |
| Ethyl acetate | 141-78-6 | Plant volatiles | Aladdin Biochemical Technology Co., LTD | Shanghai, China | 99.5 % | 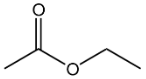 |
| Piperitone | 89-81-6 | Plant volatiles | Macklin Biochemical Technology Co., LTD | Shanghai, China | 98.0 % | 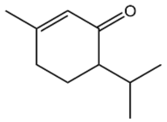 |
| Nonanal | 124-19-6 | Plant volatiles | Sigma-Aldrich LLC. | MO, USA | 95.0 % | 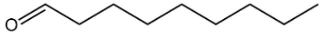 |
| Ethyl cinnamate | 103-36-6 | Plant volatiles | Aladdin Biochemical Technology Co., LTD | Shanghai, China | 99.0 % | 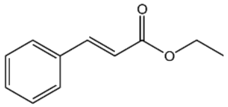 |
| *β*-Pinene | 18172-67-3 | Plant volatiles | Rhawn Biotechnology Co., LTD | Shanghai, China | 98.0 % | 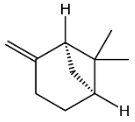 |
| Trans-2-Hexenal | 6728-26-3 | Plant volatiles | Bidepharm Medical Technology Co., LTD | Shanghai, China | 98.0 % | 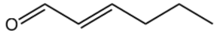 |
| Benzyl alcohol | 100-51-6 | Plant volatiles | Rhawn Biotechnology Co., LTD | Shanghai, China | 99.0 % | 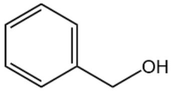 |
| 2-Phenylethanol | 60-12-8 | Plant volatiles | Aladdin Biochemical Technology Co., LTD | Shanghai, China | 99.0 % | 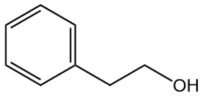 |
| Phenethyl acetate | 103-45-7 | Plant volatiles | Rhawn Biotechnology Co., LTD | Shanghai, China | 99.0 % | 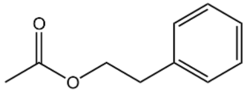 |
| cis-3-Hexenyl acetate | 3681-71-8 | Plant volatiles | Rhawn Biotechnology Co., LTD | Shanghai, China | 98.0 % | 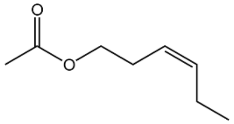 |

| Compound | Cas No. | Source | Company | Country | Purity | Structure |
|---|---|---|---|---|---|---|
| $\alpha$-Caryophyllene | 6753-98-6 | Plant volatiles | Aladdin Biochemical Technology Co., LTD | Shanghai, China | 99.0% | 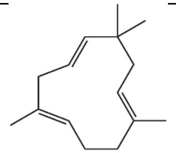 |
| Limonene | 138-86-3 | Plant volatiles | Rhawn Biotechnology Co., LTD | Shanghai, China | 95.0% | 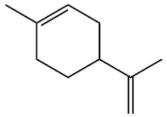 |
| Hexanal | 66-25-1 | Plant volatiles | Aladdin Biochemical Technology Co., LTD | Shanghai, China | 99.0% | 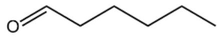 |
| Dihydro-ionone | 17283-81-7 | Plant volatiles | Rhawn Biotechnology Co., LTD | Shanghai, China | 98.0% | 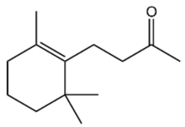 |
| 3-Octanone | 106-68-3 | Plant volatiles | Aladdin Biochemical Technology Co., LTD | Shanghai, China | 98.0% | 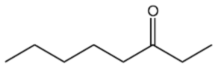 |
| 1-Octen-3-one | 4312-99-6 | Plant volatiles | Aladdin Biochemical Technology Co., LTD | Shanghai, China | 95.0% | 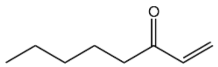 |
| 3-Octanol | 589-98-0 | Plant volatiles | Energy Chemical Technology Co., LTD | Shanghai, China | 98.0% | 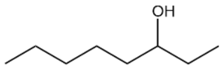 |
| 2-Methyl butyric acid | 116-53-0 | Plant volatiles | Aladdin Biochemical Technology Co., LTD | Shanghai, China | 98.0% | 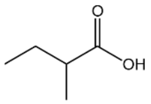 |
| $\alpha$-Pinene | 2437-95-8 | Plant volatiles | Sigma-Aldrich LLC. | MO, USA | 98.0% | 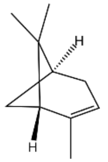 |
| Camphene | 79-92-5 | Plant volatiles | Aladdin Biochemical Technology Co., LTD | Shanghai, China | 98.0% | 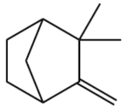 |

| Compound | Cas No. | Source | Company | Country | Purity | Structure |
|---|---|---|---|---|---|---|
| Linalool oxide | 60047-17-8 | Plant volatiles | Aladdin Biochemical Technology Co., LTD | Shanghai, China | 98.0 % | 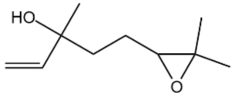  |
| (-)-Verbenone | 1196-01-6 | Plant volatiles | Aladdin Biochemical Technology Co., LTD | Shanghai, China | 95.0 % | 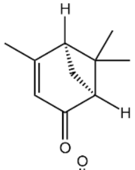  |
| Methyl benzoate | 93-58-3 | Plant volatiles | Sigma-Aldrich LLC. | MO, USA | 99.0 % | 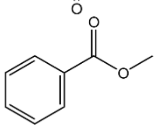  |
| M-xylene | 108-38-3 | Plant volatiles | Aladdin Biochemical Technology Co., LTD | Shanghai, China | 99.0 % | 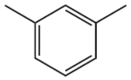 |
| Carvone | 99-49-0 | Plant volatiles | Macklin Biochemical Technology Co., LTD | Shanghai, China | 95.0 % | 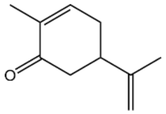  |
| 1-Octene | 111-66-0 | Plant volatiles | Macklin Biochemical Technology Co., LTD | Shanghai, China | 98.0 % | 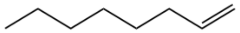 |
| 6-Methyl-5-hepten-2-one | 110-93-0 | Plant volatiles | Sigma-Aldrich LLC. | MO, USA | 99.0 % | 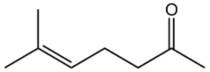  |
| Decyl aldehyde | 112-31-2 | Plant volatiles | Macklin Biochemical Technology Co., LTD | Shanghai, China | 97.0 % | 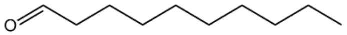  |
| Dimethyl phthalate | 131-11-3 | Plant volatiles | Aladdin Biochemical Technology Co., LTD | Shanghai, China | 99.0 % | 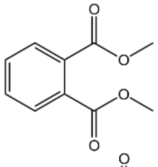  |
| Methyl cinnamate | 103-26-4 | Plant volatiles | Aladdin Biochemical Technology Co., LTD | Shanghai, China | 99.0 % | 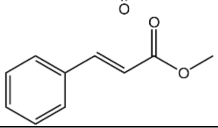  |

| Compound | Cas No. | Source | Company | Country | Purity | Structure |
|---|---|---|---|---|---|---|
| Methyl anthranilate | 134-20-3 | Plant volatiles | Aladdin Biochemical Technology Co., LTD | Shanghai, China | 99.0 % | 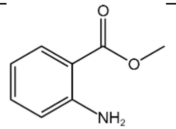 |
| *α*-Terpineol | 10482-56-1 | Plant volatiles | Aladdin Biochemical Technology Co., LTD | Shanghai, China | 98.0 % | 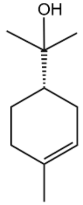 |
| (E)-*β*-farnesene | 18794-84-8 | Plant volatiles | Aladdin Biochemical Technology Co., LTD | Shanghai, China | 90.0 % | 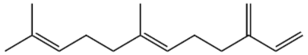 |
| Nerolidol | 7212-44-4 | Plant volatiles | Sigma-Aldrich LLC. | MO, USA | 98.0 % | 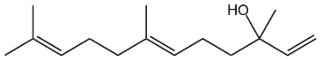 |
| Isoeugenol | 97-54-1 | Plant volatiles | Aladdin Biochemical Technology Co., LTD | Shanghai, China | 97.0 % | 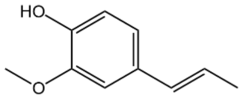 |
| Benzyl benzoate | 120-51-4 | Plant volatiles | Aladdin Biochemical Technology Co., LTD | Shanghai, China | 99.0 % | 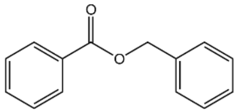 |
| 2-Hexanol | 626-93-7 | Plant volatiles | Aladdin Biochemical Technology Co., LTD | Shanghai, China | 98.0 % | 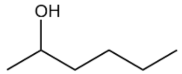 |
| Trans-3-hexen-1-ol | 544-12-7 | Plant volatiles | Bidepharm Medical Technology Co., LTD | Shanghai, China | 98.0 % | 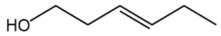 |
| *γ*-Decalactone | 706-14-9 | Plant volatiles | Aladdin Biochemical Technology Co., LTD | Shanghai, China | 98.0 % | 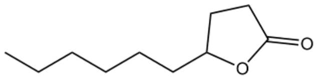 |
| Citronellyl acetate | 150-84-5 | Plant volatiles | Macklin Biochemical Technology Co., LTD | Shanghai, China | 96.0 % | 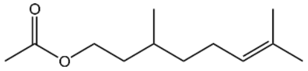 |

| Compound | Cas No. | Source | Company | Country | Purity | Structure |
|---|---|---|---|---|---|---|
| (+)-Aromadendrene | 489-39-4 | Plant volatiles | Sigma-Aldrich LLC. | MO, USA | 97.0 % | 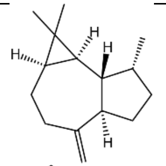 |
| Rose oxide | 16409-43-1 | Plant volatiles | Heowns Chemical TecHhnology Co., LTD | Wuhan, China | 96.0 % | 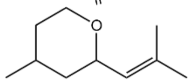 |
| 2-Tridecanone | 593-08-8 | Plant volatiles | Rhawn Biotechnology Co., LTD | Shanghai, China | 98.0 % | 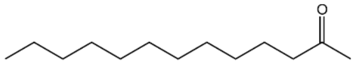 |
| Benzoylformic acid | 611-73-4 | Plant volatiles | Aladdin Biochemical Technology Co., LTD | Shanghai, China | 95.0 % | 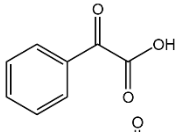 |
| Methyl anisate | 121-98-2 | Plant volatiles | Aladdin Biochemical Technology Co., LTD | Shanghai, China | 99.0 % | 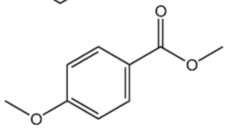 |
| Ethyl 4-methoxybenzoate | 94-30-4 | Plant volatiles | Aladdin Biochemical Technology Co., LTD | Shanghai, China | 98.0 % | 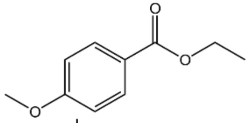 |
| $\alpha$-Phellandrene | 99-83-2 | Plant volatiles | Aladdin Biochemical Technology Co., LTD | Shanghai, China | 99.0 % | 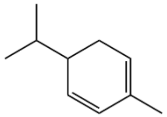 |
| Methyl acetate | 79-20-9 | Plant volatiles | Aladdin Biochemical Technology Co., LTD | Shanghai, China | 98.0 % | 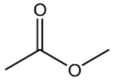 |
| Methyl decanoate | 110-42-9 | Plant volatiles | Aladdin Biochemical Technology Co., LTD | Shanghai, China | 98.0 % | 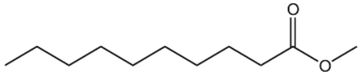 |
| $\alpha$-Ionol | 25312-34-9 | Plant volatiles | Aladdin Biochemical Technology Co., LTD | Shanghai, China | 90.0 % | 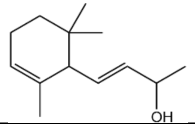 |

| Compound | Cas No. | Source | Company | Country | Purity | Structure |
|---|---|---|---|---|---|---|
| Ethyl caprate | 110-38-3 | Plant volatiles | Aladdin Biochemical Technology Co., LTD | Shanghai, China | 99.0 % | 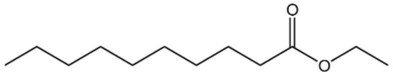 |
| Pentamethylbenzene | 700-12-9 | Plant volatiles | Bidepharm Medical Technology Co., LTD | Shanghai, China | 98.0 % | 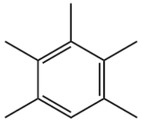 |
| Cyclohexanone | 108-94-1 | Plant volatiles | Aladdin Biochemical Technology Co., LTD | Shanghai, China | 99.0 % | 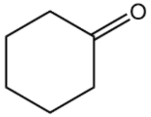 |
| *p*-Tolualdehyde | 104-87-0 | Plant volatiles | Macklin Biochemical Technology Co., LTD | Shanghai, China | 97.0 % | 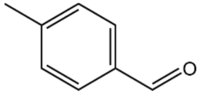 |
| Methyl eugenol | 93-15-2 | Plant volatiles | Rhawn Biotechnology Co., LTD | Shanghai, China | 98.0 % | 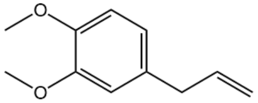 |
| trans-2-Hexenyl acetate | 2497-18-9 | Plant volatiles | Sigma-Aldrich LLC. | MO, USA | 98.0 % | 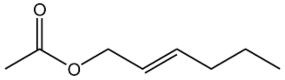 |
| 3,5-Dimethoxytoluene | 4179-19-5 | Plant volatiles | Bidepharm Medical Technology Co., LTD | Shanghai, China | 99.9 % | 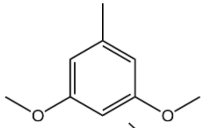 |
| Methyl jasmonate | 1211-29-6 | Plant volatiles | Sigma-Aldrich LLC. | MO, USA | 95.0 % | 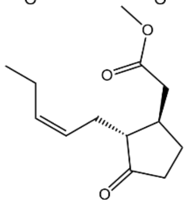 |
| 1,4-Dimethoxybenzene | 150-78-7 | Plant volatiles | Aladdin Biochemical Technology Co., LTD | Shanghai, China | 99.0 % | 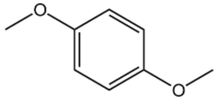 |
| Ethyl benzoate | 93-89-0 | Plant volatiles | Rhawn Biotechnology Co., LTD | Shanghai, China | 99.5 % | 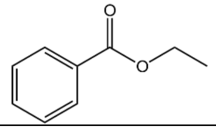 |

| **Compound** | **Cas No.** | **Source** | **Company** | **Country** | **Purity** | **Structure** |
|---|---|---|---|---|---|---|
| Bornyl acetate | 76-49-3 | Plant volatiles | Macklin Biochemical Technology Co., LTD | Shanghai, China | 97.0 % | 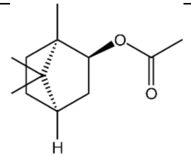 |
| Vanillin | 121-33-5 | Plant volatiles | Aladdin Biochemical Technology Co., LTD | Shanghai, China | 99.0 % | 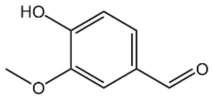 |
| *α*-Terpinene | 99-86-5 | Plant volatiles | Aladdin Biochemical Technology Co., LTD | Shanghai, China | 90.0 % | 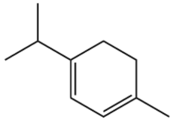 |
| Acetophenone | 98-86-2 | Plant volatiles | Aladdin Biochemical Technology Co., LTD | Shanghai, China | 99.5 % | 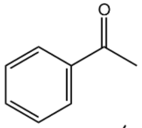 |
| Isophorone | 78-59-1 | Plant volatiles | Aladdin Biochemical Technology Co., LTD | Shanghai, China | 97.0 % | 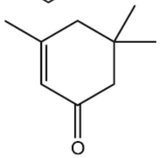 |
| *p*-Anisaldehyde | 123-11-5 | Plant volatiles | Aladdin Biochemical Technology Co., LTD | Shanghai, China | 98.0 % | 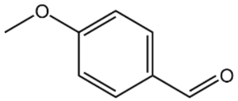 |
| trans-Anethole | 4180-23-8 | Plant volatiles | Aladdin Biochemical Technology Co., LTD | Shanghai, China | 99.0 % | 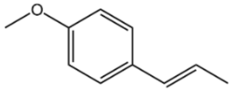 |
| cis-Anethole | 104-46-1 | Plant volatiles | Aladdin Biochemical Technology Co., LTD | Shanghai, China | 99.8 % | 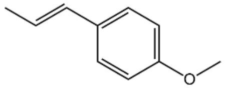 |
| *p*-Cymene | 99-87-6 | Plant volatiles | Macklin Biochemical Technology Co., LTD | Shanghai, China | 98.0 % | 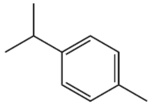 |
| (E,E)-Farnesol | 106-28-5 | Plant volatiles | Aladdin Biochemical Technology Co., LTD | Shanghai, China | 97.6 % | 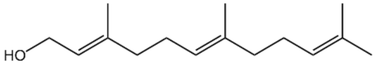 |

| Compound | Cas No. | Source | Company | Country | Purity | Structure |
|---|---|---|---|---|---|---|
| (E,E)-$\alpha$-Farnesene | 502-61-4 | Plant volatiles | Sigma-Aldrich LLC. | MO, USA | 98.0 % | 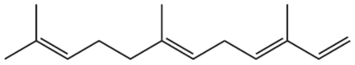 |
| 4-Allylanisole | 140-67-0 | Plant volatiles | Aladdin Biochemical Technology Co., LTD | Shanghai, China | 97.0 % | 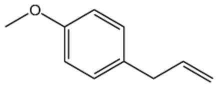 |
| 4-Oxoisophorone | 1125-21-9 | Plant volatiles | Energy Chemical Technology Co., LTD | Shanghai, China | 98.0 % | 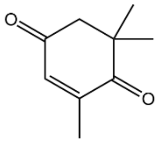 |
| 2-Aminobenzaldehyde | 529-23-7 | Plant volatiles | Bidepharm Medical Technology Co., LTD | Shanghai, China | 98.0 % | 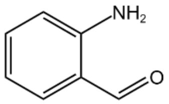 |
| 3,7-Dimethylocta-1,6-dien-3-yl acetate | 115-95-7 | Plant volatiles | Bidepharm Medical Technology Co., LTD | Shanghai, China | 98.0 % | 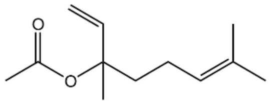 |
| 2-Methylbutyraldehyde | 96-17-3 | Plant volatiles | Bidepharm Medical Technology Co., LTD | Shanghai, China | 95.0 % | 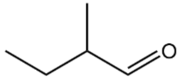 |
| 1,4-Xylene | 106-42-3 | Plant volatiles | Heowns Chemical TecHhnology Co., LTD | Wuhan, China | 98.0 % | 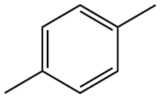 |
| Menthone | 10458-14-7 | Plant volatiles | Bidepharm Medical Technology Co., LTD | Shanghai, China | 98.0 % | 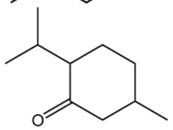 |
| 2-Nonanone | 821-55-6 | Plant volatiles | Aladdin Biochemical Technology Co., LTD | Shanghai, China | 99.0 % | 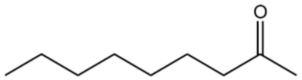 |
| $\gamma$-Octanoic lactone | 104-50-7 | Plant volatiles | Bidepharm Medical Technology Co., LTD | Shanghai, China | 98.0 % | 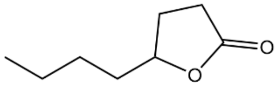 |

| Compound | Cas No. | Source | Company | Country | Purity | Structure |
|---|---|---|---|---|---|---|
| Ethyl 2-hydroxybenzoate | 118-61-6 | Plant volatiles | Bidepharm Medical Technology Co., LTD | Shanghai, China | 99.8 % | 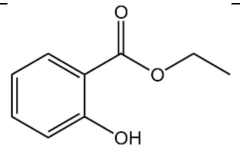 |
| Isovaleraldehyde | 590-86-3 | Plant volatiles | Aladdin Biochemical Technology Co., LTD | Shanghai, China | 98.0 % | 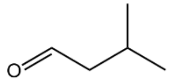 |
| 1-Decene | 872-05-9 | Plant volatiles | Aladdin Biochemical Technology Co., LTD | Shanghai, China | 95.0 % | 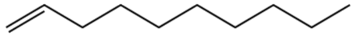 |
| 2-Pentadecanone | 2345-28-0 | Plant volatiles | Sigma-Aldrich LLC. | MO, USA | 98.5 % | 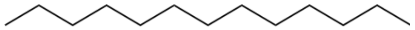 |

Table S3 Quantitative GC-MS analysis of 9-ODA and HOB.

| **Caste** | **9-ODA** | **HOB** |
|---|---|---|
| Virgin queen | 15.951 ± 6.520 μg | - |
| Mated queen | 35.993 ± 3.560 μg | 8.019 ± 1.589 μg |
| Drone | - | - |
| Worker | - | - |

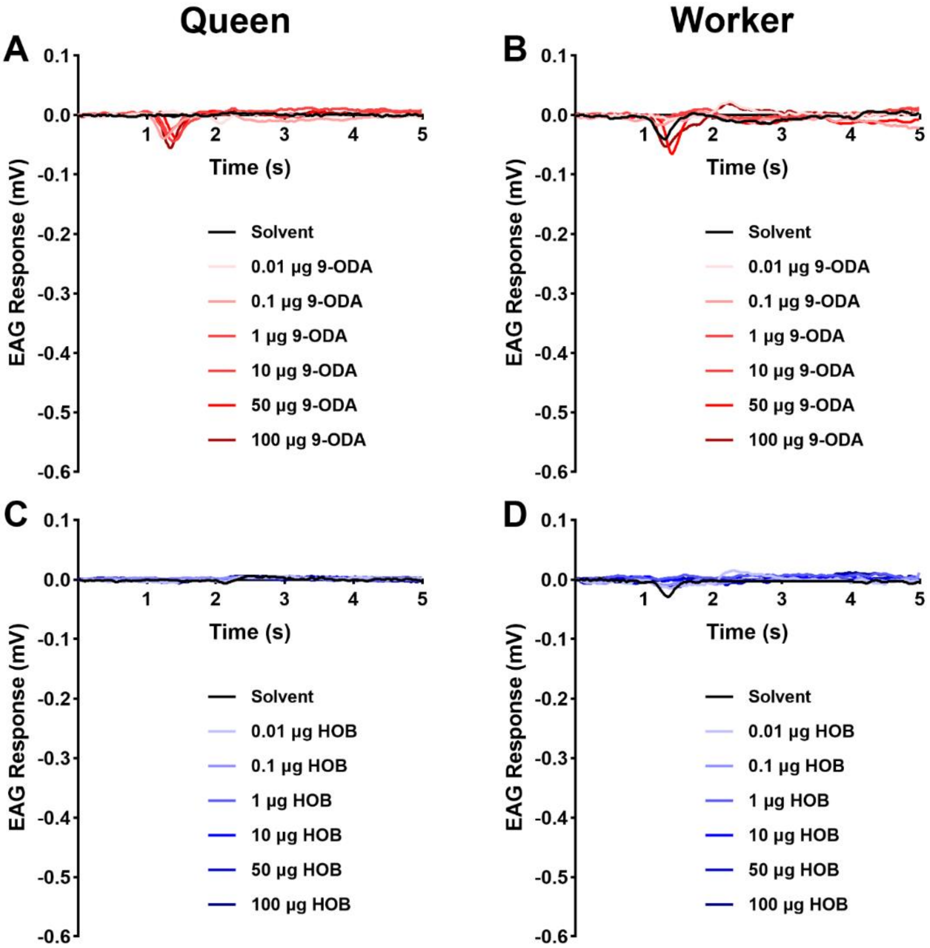


**Figure S1. EAG traces of queens and workers in response to 9-ODA and HOB.** Responses of queens and workers elicited by the increasing doses (0.01 to 100 µg) of (A, B) 9-ODA and (C, D) HOB.

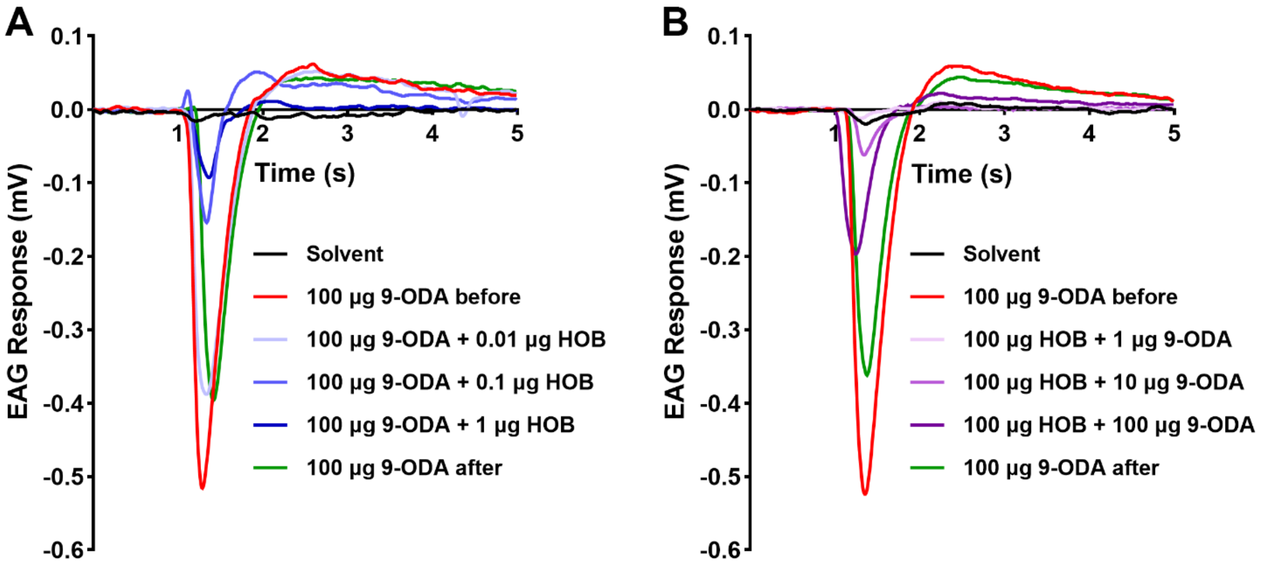


**Figure S2. Representative EAG inhibitory trace of HOB to 9-ODA.** EAG responses of (A) 100 µg 9-ODA with 0.01 to 1 µg HOB and (B) 100 µg HOB with 1 to 100 µg 9-ODA.

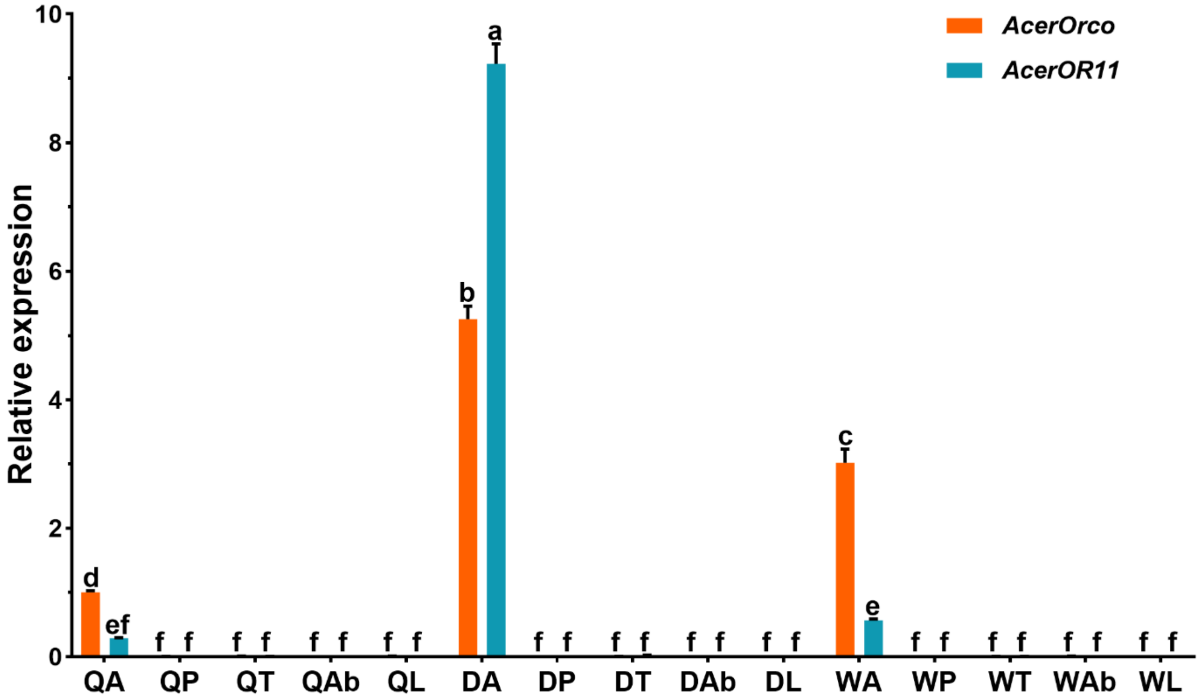


**Figure S3. RT-qPCR results of *AcerOr11* and *AcerOrco* in *A. cerana*.** QA: Queen antenna; QP: Queen proboscis; QT: Queen thorax; QAb: Queen abdomen; QL: Queen legs; DA: Drone antenna; DP: Drone proboscis; DT: Drone thorax; DAb: Drone abdomen; DL: Drone legs; WA: Worker antenna; WP: Worker proboscis; WT: Worker thorax; WAb: Worker abdomen; WL: Worker legs. Different lowercase letters indicate significant differences based on a one-way ANOVA followed by Tukey's multiple comparison test ($P < 0.05$, N=3). Data are the mean ± standard error.

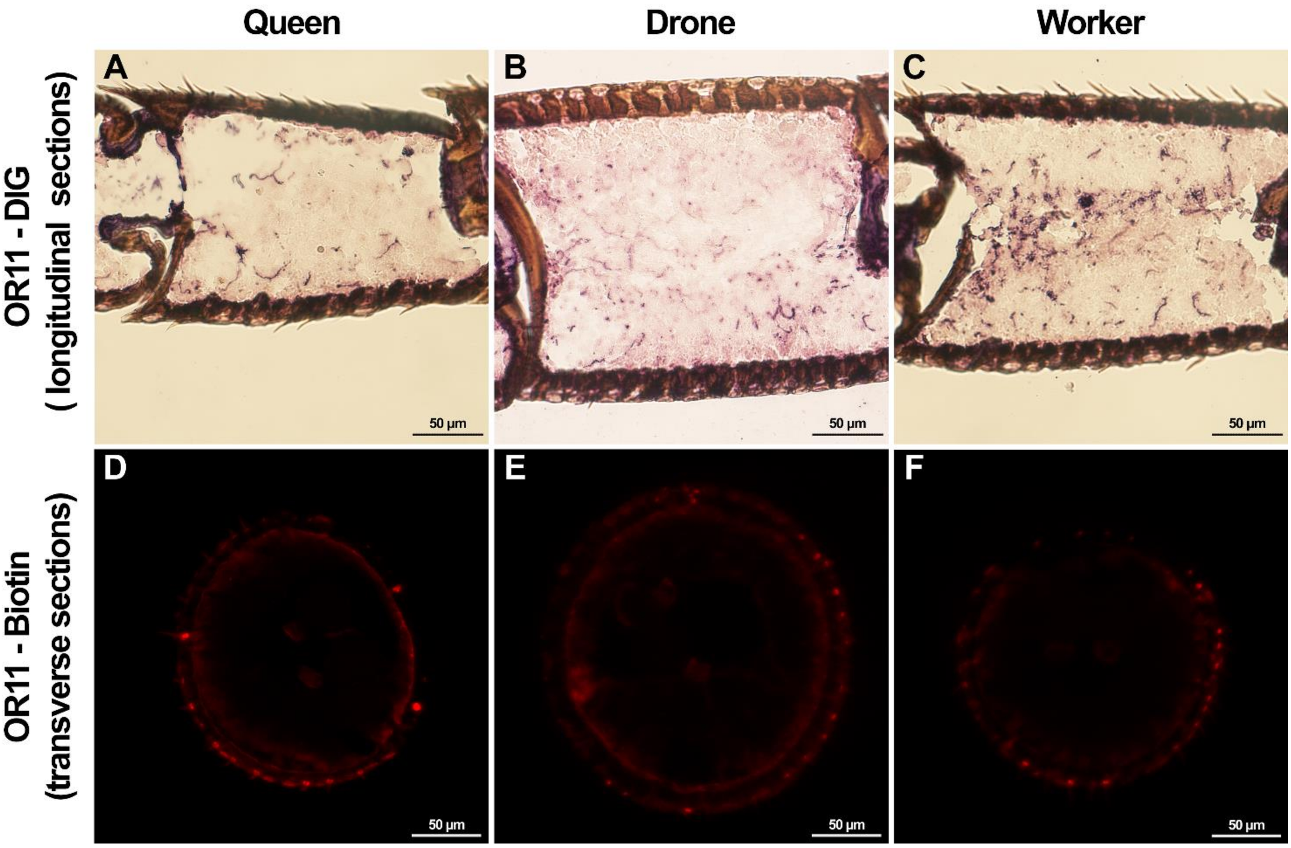


**Figure S4. Sense RNA probe of FISH of AcerOr11 from the antennae of *A. cerana*.** (A-C) In situ hybridization with the sense Or11-DIG probe on longitudinal sections. (D-F) Fluorescence in situ hybridization with the sense Or11-Biotin specific probe on longitudinal sections.

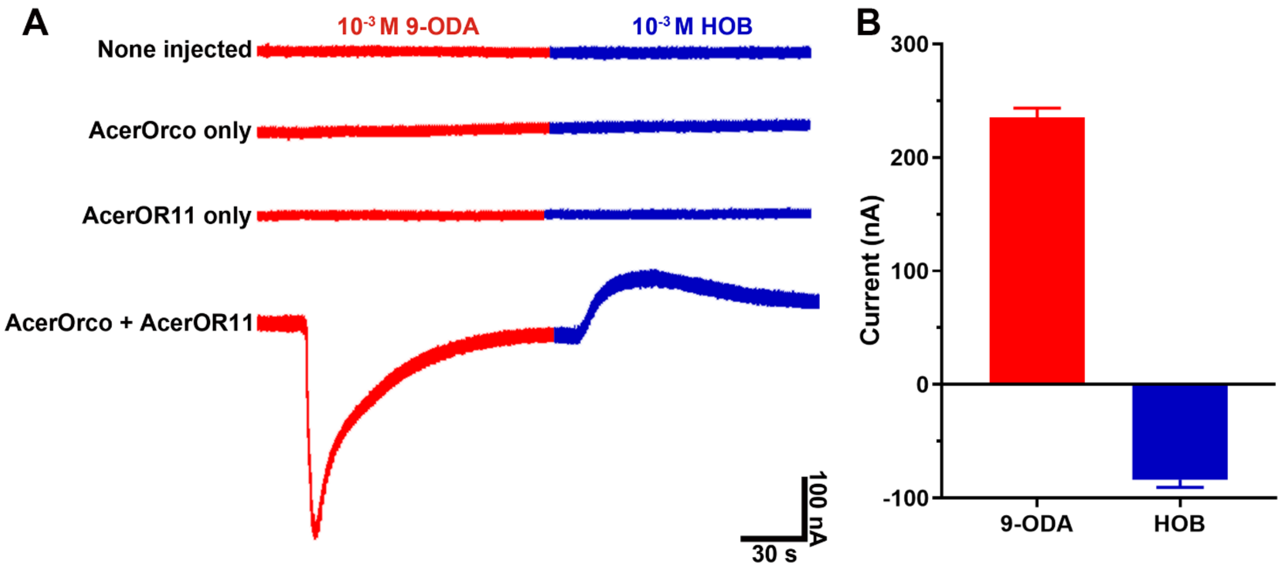


**Figure S5. Representative TEVC trace of 9-ODA and HOB.** (A) Control experiments with non-injected oocytes and oocytes expressing AcerOrco, AcerOr11, or AcerOr11/AcerOrco. (B) AcerOr11/AcerOrco expressing oocytes response to 9-ODA and HOB at $10^{-3}$ M concentration (N = 5).

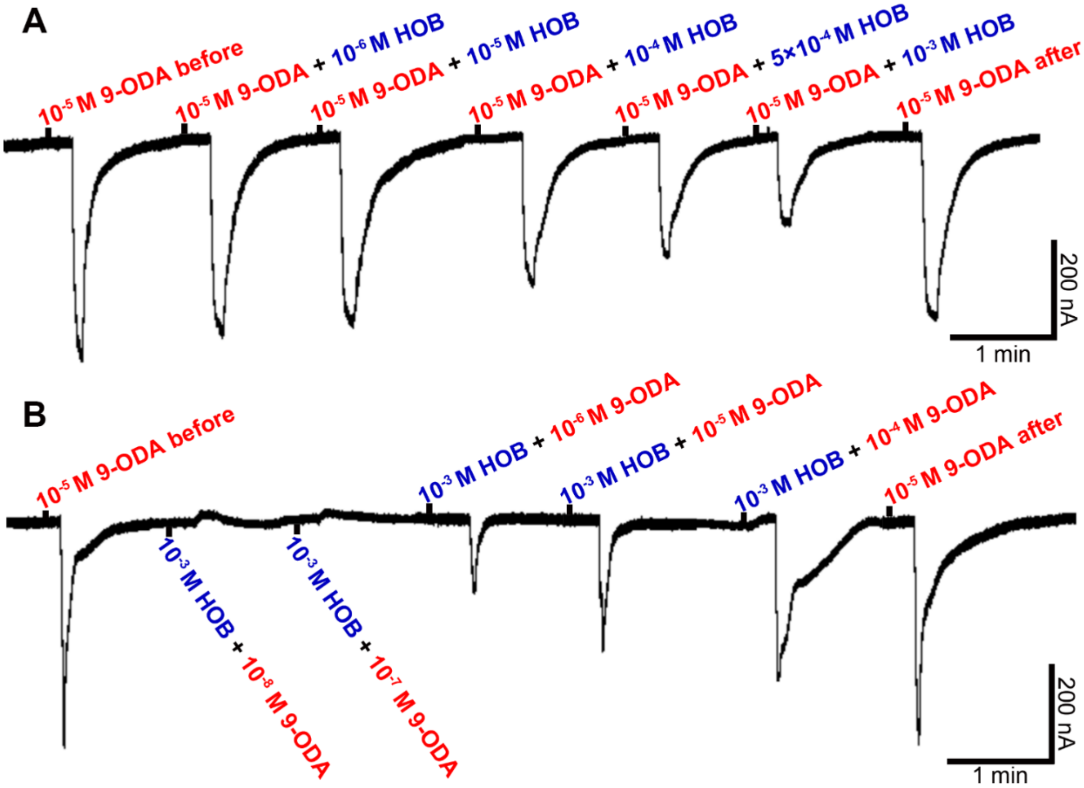


**Figure S6. Inhibitory trace of HOB to 9-ODA in *Xenopus* oocytes.** (A) TEVC response of AcerOR11 to $10^{-5}$ M 9-ODA with $10^{-6}$ to $10^{-3}$ M HOB. (B) TEVC response of AcerOr11 to $10^{-3}$ M HOB with $10^{-8}$ to $10^{-4}$ M 9-ODA.
